# *MVsim*: a toolset for quantifying and designing multivalent interactions

**DOI:** 10.1101/2021.08.01.454686

**Authors:** Bence Bruncsics, Wesley J. Errington, Casim A. Sarkar

## Abstract

Arising through multiple binding elements, multivalency can specify the avidity, duration, cooperativity, and selectivity of biomolecular interactions, but quantitative prediction and design of these properties has remained challenging. Here we present *MVsim*, an application suite built around a configurational network model of multivalency to facilitate the quantification, design, and mechanistic evaluation of multivalent binding phenomena through a simple graphical user interface. To demonstrate the utility and versatility of *MVsim*, we first show that both monospecific and multispecific multivalent ligand-receptor interactions, with their noncanonical binding kinetics, can be accurately simulated. We then quantitatively predict the ultrasensitivity and performance of multivalent-encoded protein logic gates, evaluate the inherent programmability of multispecificity for selective receptor targeting, and extract rate constants of conformational switching for the SARS-CoV-2 spike protein and model its binding to ACE2 as well as multivalent inhibitors of this interaction. *MVsim* is freely available at https://sarkarlab.github.io/MVsim/.

## Main

Multivalent interactions are fundamental building blocks of supramolecular systems. Deriving from multiple binding elements within sets of interacting molecules, multivalency is used to regulate intracellular compartmentalization^1-5^, high-avidity interactions^6-9^, ultrasensitivity^10^, and dynamics and selectivity of molecular recognition^11-13^. The expansive utility of multivalency has driven multiple computational approaches to describe aspects of multivalent interactions^14-22^. However, as the molecular systems of interest and synthetic design ambitions become increasingly manifold and complex, frameworks are required that are extensible across the multi-parameter landscape and that can be furthered into interactive tools for design and quantification. We previously developed a conceptualization of multivalency that described the noncanonical signatures of multivalent receptor-ligand interactions as the flux through an interconnected network of configurational microstates^23^. This approach provided highly-resolved mechanistic insights into the dynamical events that underlie simple multivalent interactions and indicated a means with which to extend existing experimental techniques – such as surface plasmon resonance (SPR) – to macromolecular systems previously beyond the scope of quantitative analysis due to their complexity and heterogeneity^24,25^. However, this conceptual framework was limited to monospecific multivalent interactions between proteins of certain topologies and was also not practically implementable to quickly analyze and design a wide range of molecular systems^26-29^.

Here, we have developed a new and expanded computational method based on the original conceptual framework, which we present as *MVsim*, an interactive toolset with a simple graphical user interface (GUI) for the design, prediction, multi-dimensional parameter exploration, and quantification of multivalent binding phenomena. *MVsim* enables users to simulate multivalent binding through an expansive implementation of configurational multivalent networks within the MATLAB software environment^30^. With user-specified kinetic, topological, and structural parameters, *MVsim* simulates the conformational dynamics and binding responses for multicomponent systems of multidomain, multispecific, and multimeric interacting biomolecules. Further, *MVsim* synthesizes its outputs as sets of interactive kinetic traces to facilitate visualization, inspection, quantification, curve fitting, and experimental implementation of the structure-activity relationships and the information-coding intrinsic to multivalency.

We first validate the ability of *MVsim* to accurately simulate both monospecific and multispecific protein-protein interactions, the latter of which was not possible in our initial model of multivalency^23^. To then demonstrate application of *MVsim*, we use experiment-guided modeling to quantify switch-like signaling of synthetically-designed systems^31^, uncover design rules and predict the response dynamics of multivalent logic gates^31^, and leverage multispecific ligand architectures to enable selective receptor targeting for therapeutic development^10^. Further, we apply *MVsim* toward the inspection and quantification of viral spike protein dynamics. At present, nowhere is the importance of multivalency more assertively illustrated than with the mechanics of infection and therapeutic targeting of the configurationally dynamic, trimeric SARS-CoV-2 S protein^32,33^, its dimeric ACE2 receptor^34,35^, and a growing library of designed multivalent and multispecific neutralizing inhibitors^36-40^. Here, we use *MVsim* to derive an effective concentration for the ACE2 interaction, quantify intramolecular rate constants of SARS-CoV-2 S protein receptor binding domain (RBD) conformational switching that enable host cell engagement, and probe the consequences of variants with altered conformational stability of the S protein. This series of multivalent and multi-ligand simulations served as an intuitive means to quantify the relationship between macromolecular topology and SARS-CoV-2 S protein response dynamics, infectivity, and refined approaches for therapeutic targeting.

In sum, *MVsim* offers an intuitive and easy-to-implement molecular design toolset, bringing enhanced quantification and predictive design of multicomponent and multivalent systems to protein engineering, molecular and cellular systems analysis, and therapeutic design.

## Results

### Development of *MVsim*

*MVsim is* a multivalent interaction toolset built upon our configurational microstate network model^23^, which expanded upon prior modeling efforts in the literature by explicitly treating multivalency as a dynamic ensemble of binding configurations driven through local, topology-derived effective concentrations. *MVsim* represents a reconceptualization and application of the initial, more limited network model to now provide mechanistic descriptions of an array of biologically and therapeutically relevant multivalent systems and to quantitatively predict binding responses and conformational dynamics across a breadth of parameter space^31,37,40^.

The creation of the *MVsim* toolset translated the fundamental concepts of the network model into MATLAB coding environment through a series of implementations that represent significant advances in its ability to easily simulate a broad range of multivalent interactions. First, to describe a user-specified multi-ligand, multivalent interaction system, we developed a rule-based modeling routine to automatically enumerate all possible binding microstates and configuration transitions between them to generate a descriptive kinetic rate model (Extended Methods, Supplementary Information). For example, extended to its furthest, *MVsim* simulates competitive interaction among three topologically-varied trivalent ligands for a trivalent receptor, described with a system of 1,538 differential rate equations.

Second, *MVsim* effectively and rapidly parameterizes the system of rate equations with computed topology-derived first-order rate constants of association. Here, *MVsim* uses dimensionally-reduced polar coordinate integrations of the molecular interaction volumes. With this approach, the frequency of all pairwise combinations of multivalent interaction between a ligand and receptor binding domain are calculated with joint probability density functions to yield a set of effective concentrations. This routine enables efficient calculations to be performed with high spatial resolution for nanoscale and mesoscale multivalent species with domain diameters, linkages, and persistence lengths exceeding 1000 Å.

Third, *MVsim* has an extensive multiparameter description of the molecular multivalent landscape that allows for zero-fit prediction of the response dynamics of fully-parameterized systems where experimental multivalent data is absent. Conversely, *MVsim* enables parameter estimation for topologically under-characterized systems where multivalent binding kinetics have been measured. *MVsim* facilitates quantification of multivalent binding responses in terms of effective rate constants of association 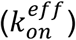 and dissociation 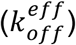, equilibrium binding constants 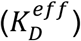, competitive inhibitor potency (*IC*_50_), and Hill coefficients describing ultrasensitive switch-like behavior.

### Parameter inputs

Interfacing *MVsim* with MATLAB’s app design environment enabled creation of a tabbed GUI to guide the specification of biologically important manifestations of multivalency via multiparameter inputs in *MVsim*. The *MVsim* GUI enables full input parameterization of the domain and linkage topologies of the ligand(s) (Fig. 1a) and receptor (Fig. 1c), the monovalent kinetics between each pairwise combination of ligand-receptor binding domains (Fig. 1a), the temporal ligand concentration dynamics (Fig. 1b), and the set of parameters that govern SPR and related kinetic studies, including the association and dissociation times, flow rate, and level of immobilized receptor. Additionally, for instances where detailed topological information is known, *MVsim* allows users to directly input effective ligand concentrations and end-to-end probability density functions for the multivalent system of interest (Fig. S1). Finally, once a multivalent design has been input, the user can initiate *MVsim* and subsequently survey a range of non-topological parameter variants in quick succession (Fig. 1d).

**Fig. 1:**
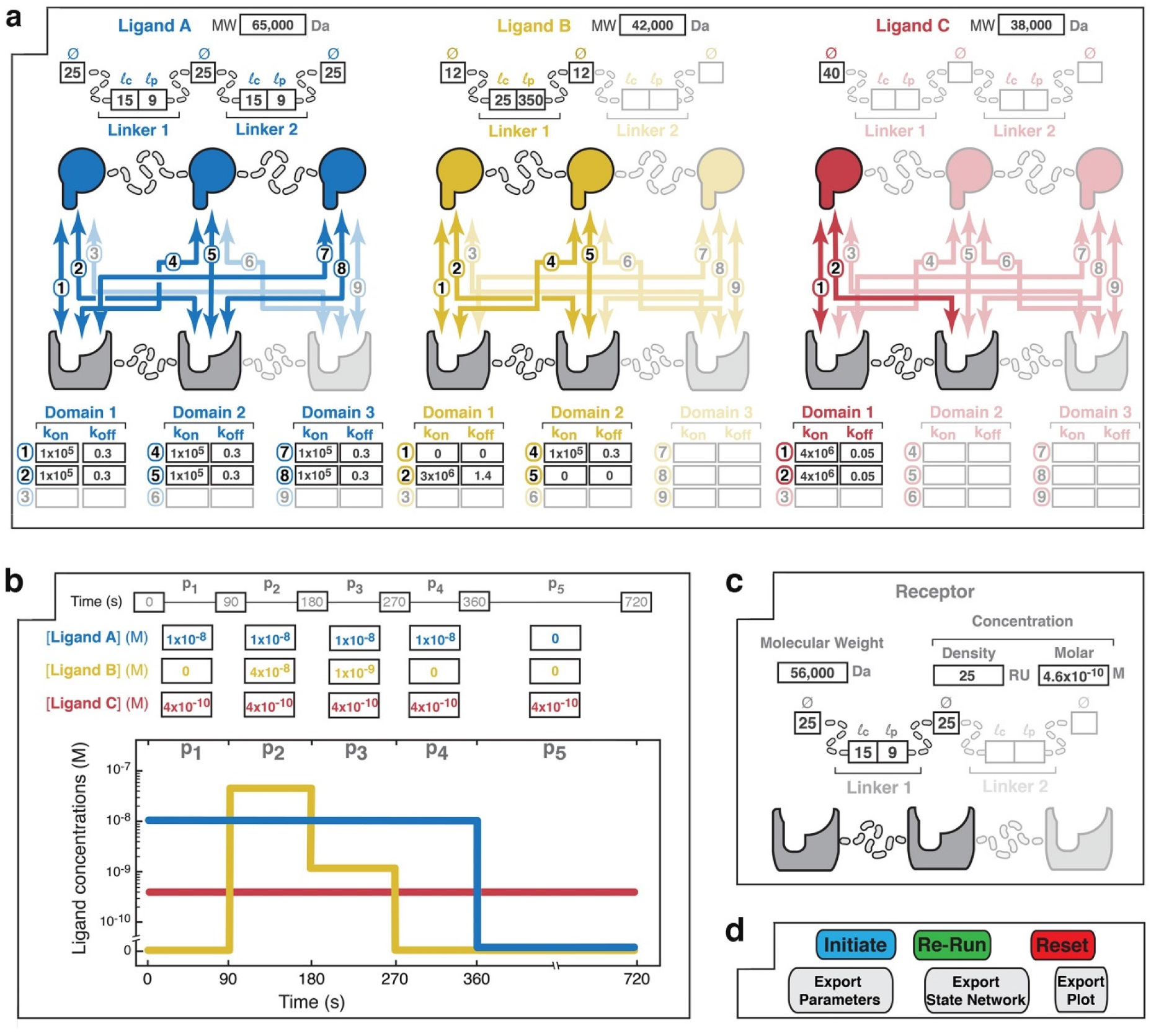
The *MVsim* input design interface provides interactive parameter specification for systems of multivalent, multi-molecular interaction. a,b,c,d. The user specifies the design of the multivalent system of interest across individual parameterization tabs within the MATLAB app environment. (**a**) A point-and-click interface enables the user to select the number of ligands (up to three) and valencies of the ligand(s) and receptor (up to trivalent) that compose the multivalent system. Based upon the chosen design, the user specifies the structure of each of the ligands by entering the applicable binding domain diameters (Ø), linker contour lengths (*l*_c_), and linker persistence lengths (*l*_p_). Further, the applicable combinatorial interactions (numbered 1 to 9) unique to each receptor-ligand pairing are highlighted. Tabulated parameter fields allow the user to input monovalent rate constants for each pairwise site of interaction in the multivalent system. Non-binding interactions can be indicated with *k*_*on*_ and *k*_*off*_ values of zero. (**b**) Below the kinetic input fields, a tabulated input field allows the user specify patterns of total, bulk ligand concentrations across specified points during the timecourse of the interaction. An association phase occurs during periods of non-zero bulk ligand concentration (e.g., 90-270 s for Ligand B). Dissociation phases occur when ligand is removed from the bulk solution (e.g., 360-720 s for Ligand A). Here, Ligand C is specified as continuously present in solution during the 720 s of the interaction timecourse. The graphical display allows visualization of the specified bulk concentration pulse pattern. Ligand pulses can be added or removed (up to 7 in total). (**c**) User input parameters for the receptor. Receptor concentration can be specified as either an SPR-mimicking surface density (measured in RU; where 1 RU equals ∼1 pg/mm2) or a molar concentration. Receptor topology is specified in the same form as described above for the ligands. (**d**) The *MVsim* controller tab enables initiation, iteration, and export of binding simulations. “Initiate” executes a user-parameterized simulation. “Re-run” executes an abbreviated and faster simulation for use where no changes were made to input parameters that alter the valency or topology of the system. “Reset” relaunches the app and clears user input parameters from all fields.

### Simulation outputs

Following the initiation of a simulation, *MVsim* provides users a variety of means to visualize, interact with, and export the simulated response kinetics. Most simply, the simulation results are displayed within the output field as an interactive plot of binding response signal as a function of the specified association and dissociation time (Fig. 2). Here, users can choose between two graphical outputs of the response kinetics. First, as is typical of experimental binding kinetic data, a plot of a user-specified ligand concentration is displayed (Fig. 2a). Alternatively, users can select a plot of all composite microstate configurations underlying the response signal (Fig. 2b), binned according to valency class (Fig. 2c) or observe the competitive binding dynamics among multiple ligands (Fig. 2d). *MVsim* additionally enables users to visualize the dynamic evolution of the microstate network via an interactive map (Fig. 2e) and to export the response kinetics as a set of tab-delineated text files to facilitate deeper analysis through offline plotting and curve fitting, and through microstate network analysis within the Cytoscape software environment^41^. *MVsim* additionally enables users to directly inspect both the computed probability density functions and effective concentrations (Fig. S2).

**Fig. 2:**
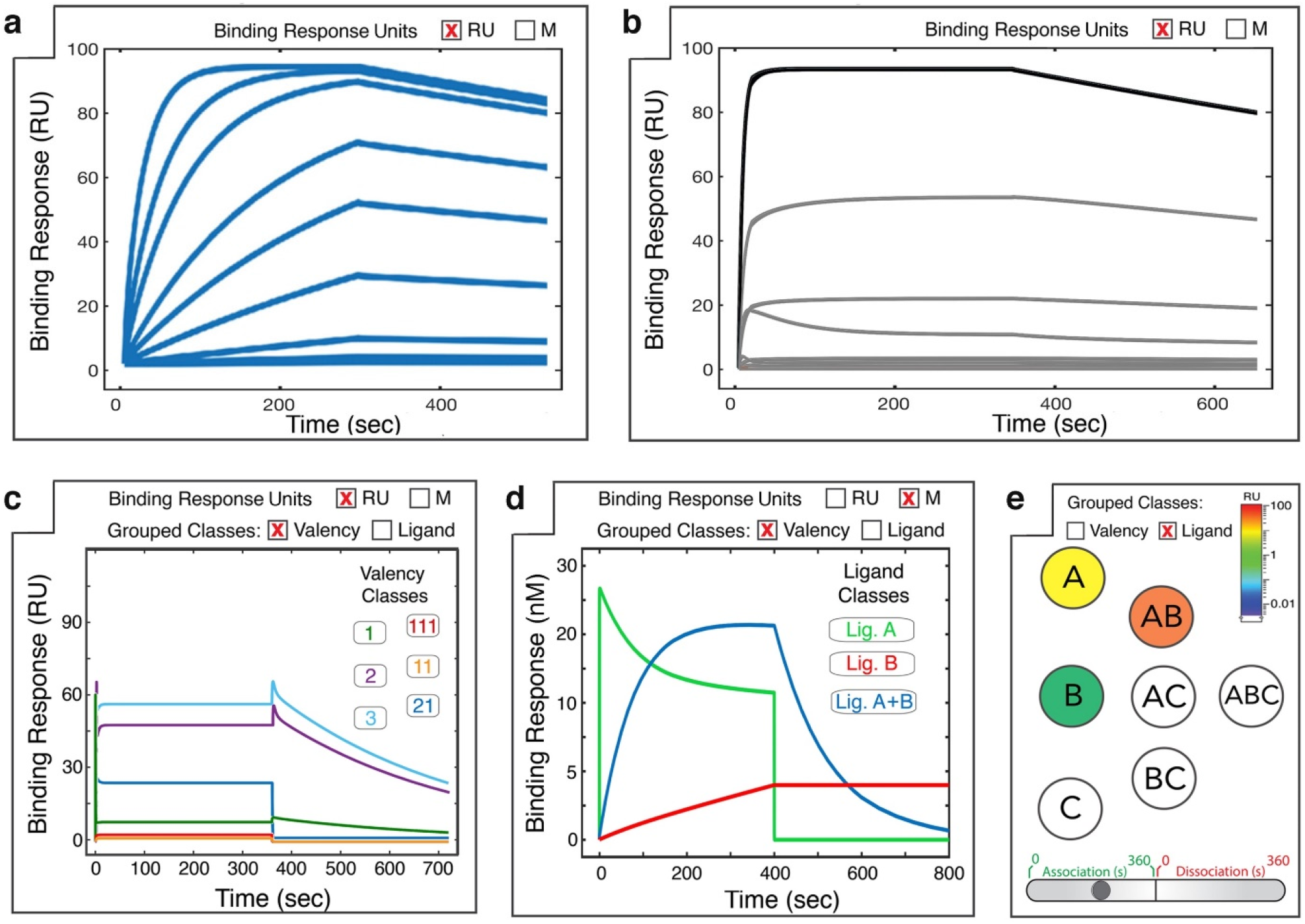
*MVsim* generates a series of outputs that enable interactive visualization of binding responses, configurational microstates, and multi-ligand dynamics. a. A simulated SPR sensorgram displays the overall response dynamics (i.e., summation of all ligands and binding microstates) for specified ligand concentration(s). Indicated here are the binding responses for a serial dilution of a single ligand binding to a receptor with association (0-300 s) and dissociation (300-600 s) phases. For simple quantitative comparison between simulations, an overall effective KD can be calculated by the equilibrium method. **b**, For a specified ligand concentration, all composite microstates are displayed. **c**,**d**, To facilitate analyses of the binding responses, the microstates can be binned according to either **(c)** valency or **(d)** ligand class. **e**, For visual analysis of the evolution of a network of microstates in (**b**), an interactive graph shows population changes in microstate classes over a timecourse of association and dissociation.

### Assessing performance against experimental model systems

To evaluate the accuracy of simulating complex topologies, *MVsim* was used to predict the binding response dynamics of our previously constructed monospecific multivalent interactions and two new experimental systems (Fig. 3). In agreement with our previous studies of monospecific interactions, *MVsim* predicts monospecific multivalent binding (Fig. 3b,c) but now also shows improved sensitivity to the topological constraints that can impede certain configurations, such as those that require contorted twisting of interdomain linkages. We further used *MVsim* to make new predictions in both multispecific ligand and multi-ligand interaction systems. In the first validation, a multispecific receptor-ligand architecture was designed using two sets of protein-protein interaction domains (Fig. 3a). Experimentally determined monovalent kinetic rate constants and structure-derived molecular topologies were used to parameterize the model (Fig. 3a,d) and generate a simulated dataset (Fig. 3e). Comparing simulation with the corresponding SPR dataset (Fig. 3f) demonstrates a good agreement with regard to the ability of *MVsim* to predict *a priori* the magnitude and multiphasic character of the experimental binding responses. Further, *MVsim* provides mechanistic explanations for these binding responses, showing, for example, the contribution of high-stoichiometric configurations to the microstate ensemble driven by the use of rigid, α-helical linkers (Fig. S3a-c).

**Fig. 3:**
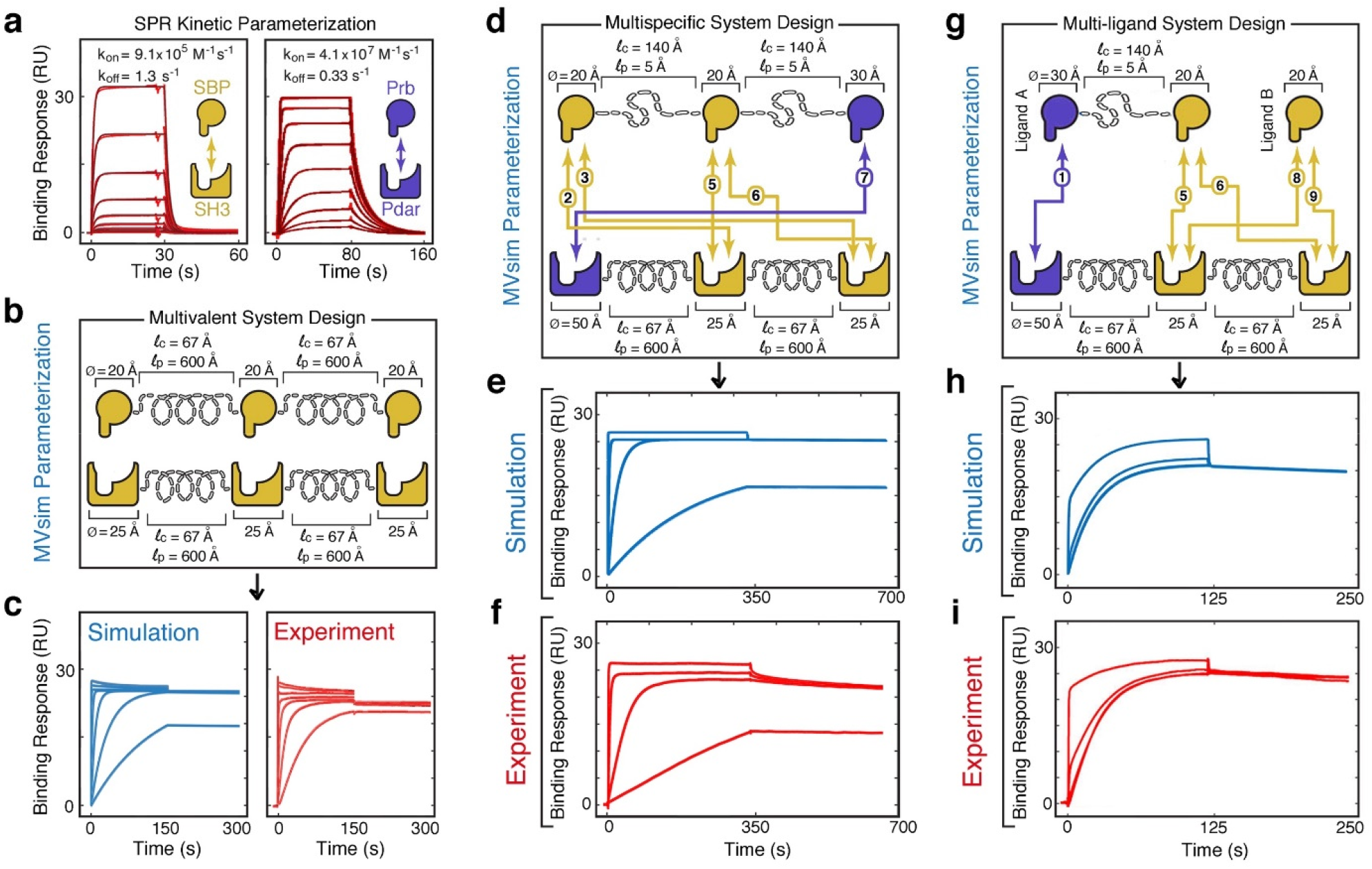
*MVsim* accurately simulates beads-on-a-string multivalent, multispecific and multi-ligand interactions. a. Monovalent SPR kinetic rate constants were experimentally determined for the SH3-binding peptide (SBP)-SH3 (**a**, left panel) and Prb-Pdar (**a**, right panel) interactions that were used to build the multivalent systems used in this study. **b**, A trivalent, monospecific receptor-ligand interaction was engineered and parameterized within *MVsim* using values for the kinetic rate constants of association (*k*_on_) and dissociation (*k*_off_), diameters (Ø) for the protein-protein binding domains, and contour (*l*_c_) and persistence (*l*_p_) lengths for the alpha helical linkers. **c**, Simulated (**c**, left panel) and experimental (**c**, right panel) binding response dynamics for the trivalent, monospecific interaction. An overlay is shown of binding responses for six simulated ligand concentrations (5, 15, 60, 250, 1000, and 2000 nM). **d**, A trivalent, bispecific receptor-ligand interaction was engineered and parameterized within *MVsim* using values for the kinetic rate constants of association (*k*_on_) and dissociation (*k*_off_), diameters (Ø) for the protein-protein binding domains, and contour (*l*_c_) and persistence (*l*_p_) lengths for the alpha-helical linkers. The bidirectional arrows indicate compatible interactions between the receptor and ligand binding domains. **e**, Simulated binding response dynamics modeled by *MVsim* for the parameterized trivalent, bispecific interaction. An overlay is shown of binding responses for four simulated ligand concentrations (5, 25, 100, and 1000 nM). **f**, Experimental SPR binding response dynamics for the trivalent, bispecific interaction at the same four ligand concentrations as in (**e**). **g**, The Pdar-Prb and SBP-SH3 protein-protein binding domains were used to create a multi-ligand system. **h**, Simulated binding response dynamics modeled by *MVsim* for the parameterized dual ligand system. An overlay is shown of binding responses for three simulated mixtures of ligands A and B (1 nM A + 2.5 nM B; 1 nM A + 50 nM B; and 1 nM A + 250 nM B). **i**, Experimental SPR binding response dynamics for the same three dual ligand mixtures as in (**h**).

As a second model-experiment validation, a fully parameterized, dual ligand interaction system was constructed (Fig. 3g). Again, comparing the simulated binding responses (Fig. 3h) to the corresponding experimental SPR data (Fig. 3i) shows good agreement between *MVsim* and experiment in the characteristic multiphasic appearance of both the association and dissociation phases of the interaction. Moreover, beyond simply predicting the overall kinetics of the system, *MVsim* provides insights into the mechanics of multiple multivalent and multispecific ligands competing for a receptor, and attributes these molecular properties back to the macroscopically observable features of the multiphasic binding responses. Here, for example, *MVsim* quantitatively captures how effective rate constants of dissociation can be dictated by valency and can be used to effect temporal ordering of interactions between a rapidly, but more transiently, binding monovalent ligand and a slower, but more avid, multivalent ligand (Fig. 3d-f).

### Applications to multivalent system design and quantification

*MVsim* was established to both guide the design and implementation of multivalent properties and to facilitate parameter estimation for existing and incompletely characterized natural and synthetic multivalent systems. Here, the model’s lack of reliance upon fitted parameters enables *MVsim* to better describe the additive, competitive, and cooperative relationships implicit between kinetic, topological and valency parameters and to apply these to the quantification of multivalent properties, such as effective concentration, avidity, and binding selectivity. To evaluate the performance of *MVsim* as a molecular design and quantification tool, we assessed its ability to design and predict binding response dynamics in four different instances and applications of multivalency.

### *MVsim* predicts ultrasensitive behavior in engineered protein switches and logic gates

The effective concentration that drives multivalent binding gives these systems the inherent ability to produce nonlinear input/output response dynamics. It has been previously demonstrated, for example, that ultrasensitive toggling can be driven through the introduction of monovalent counterparts into a multivalent system^31^. Dueber et al. showed that cooperative competitive dissociation of multivalent protein-protein complexes effects switch-like transitions that can be leveraged to control the fractional saturation of receptor-ligand interactions and enzymatic activity. Here, we apply *MVsim* to study the activation dynamics of engineered bivalent and trivalent protein switches and identify critical parameters for optimal system performance. *MVsim* quantitatively predicts the relationship between the valency of the system and the magnitude of its cooperative transition to an active state (Fig. 4a). The functional range of multivalent switches can be extended through the incorporation of multispecific interactions. This design approach enables the creation of AND logic gates in which a switch response is elicited only by a programmed combination of molecular inputs. Using *MVsim* to model this system recapitulates the experimental three-input response (Fig. 4b) and reveals potential sources of erroneous activation in the design (Fig. S4). It also identifies the importance of minimizing steric imposition between intramolecular interactions, closely matching multispecific binding strengths across domains within the multi-domain construct, and ensuring the concentrations of the monovalent agonists required to facilitate dynamic switching are compatible with applications in cells and organisms (Fig. S4).

**Fig. 4:**
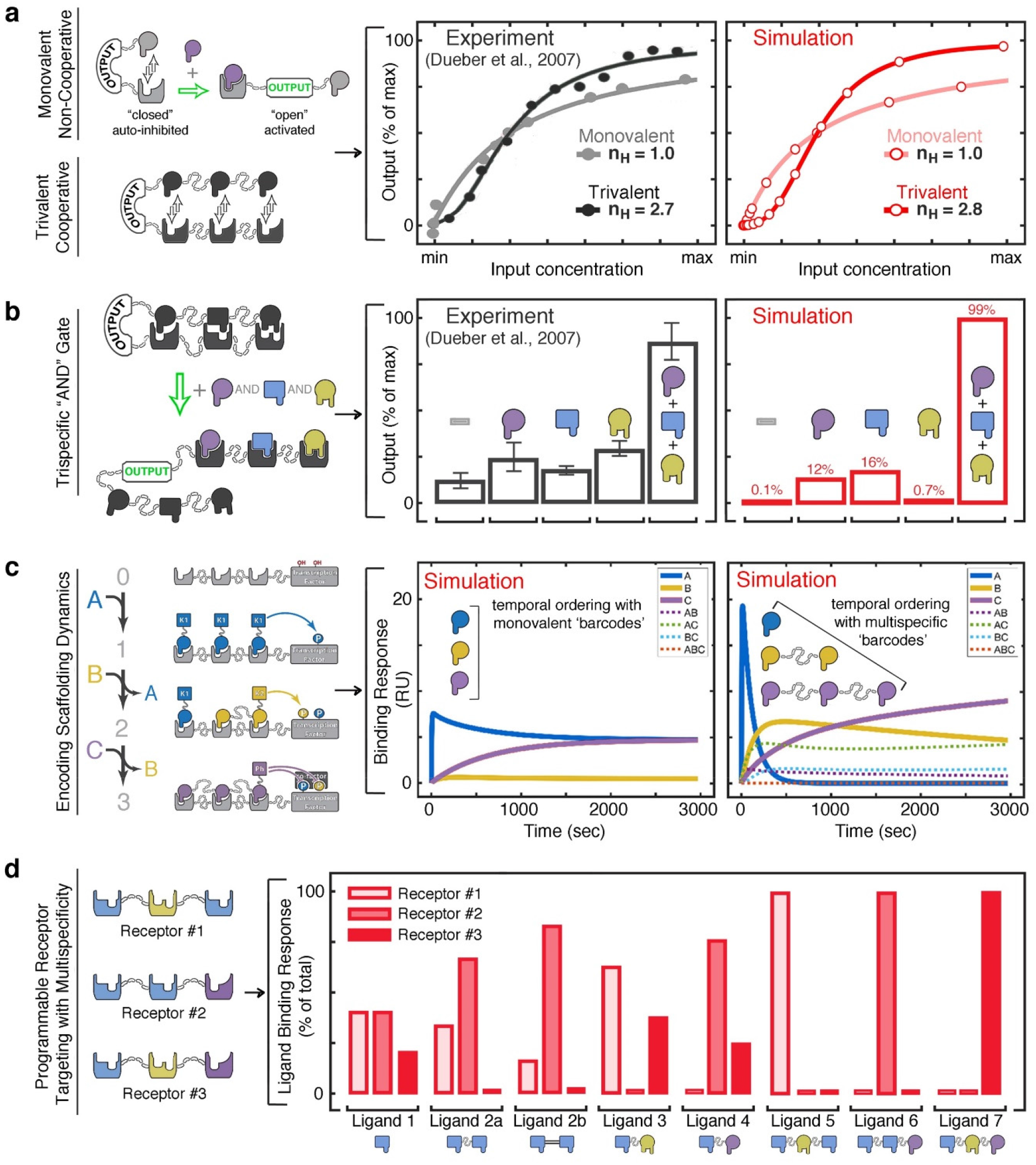
*MVsim* predicts and informs the design of switch-like dynamics, logic operations, temporal coding, and target-receptor selectivity implicit to multivalent systems. a. Experimental response dynamics of synthetic monovalent and trivalent switches from Dueber et al. were used to benchmark the predictive performance of *MVsim* simulations described by the reported structural, topological, and kinetic parameters. Ultrasensitivity of each simulated response is reported with a calculated Hill coefficient (n_H_) for direct comparison with the reported literature values. **b**, Experimental output responses for a trispecific AND logic gate, also from Dueber et al., benchmarked against an identically parameterized system in *MVsim*. For clarity, the AND gate is depicted here as inline, but the actual topology necessitates consideration of twisted configurations (detailed in Fig. S4). **c**, Parameterization of *MVsim* for the design of temporally-encoded multivalent barcodes to achieve fast and sequential interactions in a model system consisting of two kinases and a phosphatase, all of which act on a signaling hub (left schematic). Here, a simulated multispecific design enables an orderly progression of binding events (right plot) that is not accessible with monovalent counterparts (left plot). **d**, *MVsim* specifies optimal design of multivalent and multispecific ligands to yield desired patterns of selective interactions within a pool of three receptors with common binding elements.

### *MVsim* informs the use of multispecificity for molecular recognition and therapeutic targeting

Multispecificity is a potent molecular design element that is widely used in drug discovery and cell engineering. By leveraging two or more distinct binding epitopes, multispecific interactions are employed to engineer highly avid and selective molecular recognition for use in such applications as bispecific therapeutic antibodies^10,27^ and chimeric antigen receptor T cells^42^. Multi-site recognition additionally enables higher-order information processing, allowing these multispecific systems to generate differential outputs to varying combinations of inputs^11,12^.

Because the network model of multivalency computes multivalent binding as the cooperative sum of its composite interactions, *MVsim* is well-suited to the study of multipartite interactions. Here, we explore two important applications of engineered multispecificity.

First, *MVsim* examines the information-coding capacity of multispecific interactions to effect temporal ordering in a model regulatory system consisting of two kinases and a phosphatase (the ‘ligands’) that can consecutively engage a common signaling hub (the ‘receptor’) despite all three enzymes being simultaneously introduced. Here, exploration of the simulated parameter space revealed a design that enables serial binding events by exploiting the cumulative effects of concurrent binding afforded by multispecificity, the cooperative, competitive binding of multi-ligand dynamics, and the generation of effective dissociation rate constants via multivalency (Fig. 4c). Together, appending these multispecific interaction domains to the simulated regulatory system effectively creates a molecularly encoded program with biochemical and biophysical properties that specify the orderly progression of multi-ligand binding to enhancing system performance or specificity.

Second, multispecific interactions can be designed that maximally exploit any degree of variation in the type and number of surface receptors and antigens within a population for the purposes of selective targeting^10^. In this regard, we directed *MVsim* to address a design question: given a population of three distinct types of antigenic cell surfaces, what are the optimal ligand designs that can singly, doubly, and triply interrogate the population? *MVsim* demonstrates that the composition of the target receptor serves as a generally useful guide for ligand design, as seen, for example, in Fig. 4d in the relative selectivites of mono-, bi-, and trispecific Ligands 1, 3, and 7, respectively, for Receptor 3. Moreover, selective recognition can be further maximized using designed linkages that leverage the spatial proximity between receptor target surfaces; in Fig. 4d, Ligand 2b (rigid linkage) has greater selectivity than Ligand 2a (flexible linkage) for Receptor 2.

### *MVsim* models the multivalency and avidity of SARS-CoV-2 S protein interactions

At present, one of the most prominent and consequential displays of multivalent binding surrounds the surface spike (S protein) of SARS-CoV-2. The S protein is a sophisticated, conformationally activated molecular system that mediates the selective recognition of target cells and generates the driving force needed to overcome the energy barrier of membrane fusion, thus enabling viral entry into the host^32,33^. The multimeric and multivalent configuration of the S protein is central to these functions^43^. Trimeric assembly serves to stabilize the S protein against erroneous fusogenic conformational changes, establish allosteric control, and potentially present multiple receptor binding domains (RBDs) that bind multivalently with a host cell surface populated with dimeric ACE2 receptor proteins^43^. In response to these natural displays of multivalency, this same principle has been mobilized in therapeutic designs intended to neutralize, inhibit, or otherwise uncouple the structure-activity relationship of the S protein^35-40^.

Despite the significantly more complex multivalent architecture of the S protein compared with our previously described applications, *MVsim* can be effectively parameterized to model and quantify critical structural properties of the S protein-ACE2 interaction (Fig. S5a,b). For example, it remains an open question the extent to which the trimeric S protein can multivalently engage a bivalent ACE2 receptor. This question is of considerable importance for our understanding of how the affinity and avidity of S protein binding relate to infectivity, and what consequences this poses for therapeutic inhibition^40,44^. In synthetically engineered multivalent instances of the RBD-ACE2 interaction, *MVsim* quantitatively predicts the relative lack of steric hindrance that affords the ultra-high-avidity binding observed in a study by Chan et al.^40^ (Fig. 5a,b). In contrast, experiments performed on more biologically mimetic S protein-ACE2 interactions indicate a significant impediment toward high avidity binding^40^. Using *MVsim* to fit therapeutic neutralization datasets reveals an effective ligand concentration, [*L*_*eff*_], for the second engagement event between S protein and ACE2 that is 2000-fold less potent than that observed in the sterically unimpeded system (Fig. 5c,d)^40^. This inability to achieve high-avidity binding (e.g., a network in which >95% of the populated microstates are bound with maximal valency, as is the case for the ‘High’ simulation in Fig. 5d) can be explained by the combination of the rigidity of the ACE2 dimer and the apparently constrained, directional motion of the linkage tethering the RBD. Quantitative modeling approaches such as these indicate a significant potential for therapeutic designs that can potently outcompete the RBD-ACE2 interaction by leveraging multivalent binding in ways inaccessible to the S protein (Fig. 5e,f). Specifically, *MVsim* predicts that up to 1000-fold enhancements in *IC*50 values can be achieved through the use of topologically precise and constrained linkages within a designed, trivalent multispecific neutralizing therapeutic (Fig. 5). Reciprocally, *MVsim* further demonstrates how bivalency can be effectively leveraged with appropriate linkages to avidly block the RBD binding surfaces of the ACE2 dimer (Fig. S5c,d).

**Fig. 5:**
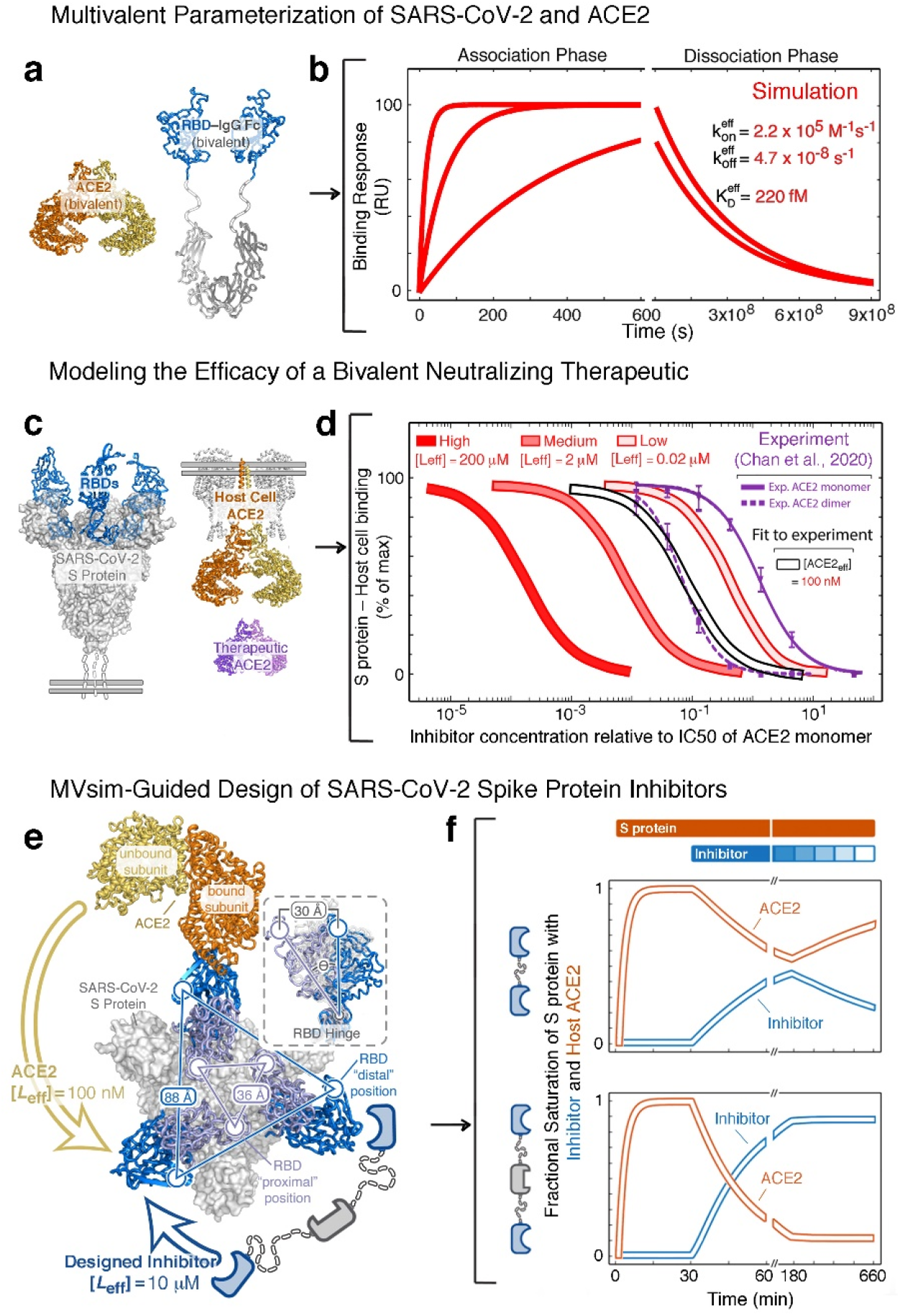
*MVsim* models the multivalency of the SARS-CoV-2 S protein RBD and ACE2 interaction and suggests novel therapeutic design strategy. a. An idealized, flexible synthetic design of an ACE2–RBD bivalent architecture. Here, the synthetic design removes the RBD from the biologically-relevant and constrained context of the rest of the S protein. **b**, The flexible RBD linkers afford a high-avidity bivalent interaction with the dimeric ACE2 that was beyond the quantification limits of the experimental SPR. Here, *MVsim* was parameterized with the features of the experimental systems and offers prediction and quantification of the ultra-high-avidity interaction. **c**, Application of *MVsim* to a biologically-relevant instance of the SARS-CoV-2 S protein RBD and ACE2 interaction. Here, the therapeutic neutralizing activity of soluble, dimeric ACE2 (purple) was quantified in a SARS-CoV-2 pseudovirus-host-cell system^40^. **d**, The resulting *IC*_50_ datasets were applied to *MVsim* in order to fit for a more biologically-relevant determination of the multivalent binding capacity of the S protein-ACE2 interaction. The experimental data (purple traces) are adapted from Chan et al.^40^ The best fit from *MVsim* gave an [*L*eff] of 100 nM (curve outlined in black), falling between the ‘Low ‘ and ‘Medium ‘ standard curves (shades of red), which represent no capacity and a modest steric capacity for bivalent binding, respectively. These simulations indicate that the RBDs in the full context of the S protein are significantly impeded for direct bivalent binding to ACE2. **e**, This steric impediment can be exploited to maximize neutralizing potency by fully leveraging therapeutic multivalency. **f**, *MVsim* can test design of neutralizing inhibitors that maximally outcompete the ACE2 interaction. Designs leveraging monospecific bivalency (top panel) and trivalent bispecificity (bottom panel) are computationally modeled for their neutralizing strengths and off-rate dependent pharmacokinetic half-lives in the presence of constant S protein (orange bar above plot) and decaying concentration of inhibitor (blue bar).

### *MVsim* quantifies the dynamics of SARS-CoV-2 S protein conformational switching

In addition to the sterically impeding immobility of the S protein-ACE2 interaction, multivalent engagement is limited by the accessibility of the RBDs as they dynamically sample configurations ranging from the occluded yet stabilized “RBD-down” conformation to the labile yet ACE2-binding competent “RBD-up” conformation^33,43^. The dynamics of this range of RBD motion are a significant target of selective pressure as the benefits of maximizing host-cell binding are countered by the need to stabilize the S protein against spontaneous fusogenic conformational change and immune surveillance of exposed critical surfaces^43^. To examine intramolecular conformational changes that yield multivalent binding, we applied *MVsim* to simulate a multicomponent experimental system consisting of a stabilized trivalent S-protein, a set of first-order rate constants *k*_*up*_ and *k*_*down*_ describing RBD conformational change, and a trivalent ligand specific for the RBD-down conformation (Fig. 6a). *MVsim*, constructed and parameterized in this way, succeeded not only in recapitulating the experimental multiphasic kinetic traces obtained in a study by Schoof et al.^37^ (Fig. 6b,c), but also in relating these response dynamics to the rates of RBD conformational switching. This modeling routine allowed *MVsim* to be used to extract unique parameter values for this conformational switching: *k*_*up*_ = 0.011 s^-1^ and *k*_*down*_ = 0.006 s^-1^ (Fig. 6d). These correspond to individual RBD half-lives of ∼2 min in the RBD-up configuration and ∼1 min in the RBD-down configuration for this stabilized, *in vitro* S protein system. To assess the accuracy of the *MVsim*-derived values of *k*_*up*_ and *k*_*down*_, these rate constants were used to parameterize a simulated SPR experiment probing the lifetime of the stabilized RBD-all-up state. Here, good agreement was observed between model and experiment (Fig. 6e,f)^37^.

**Fig. 6:**
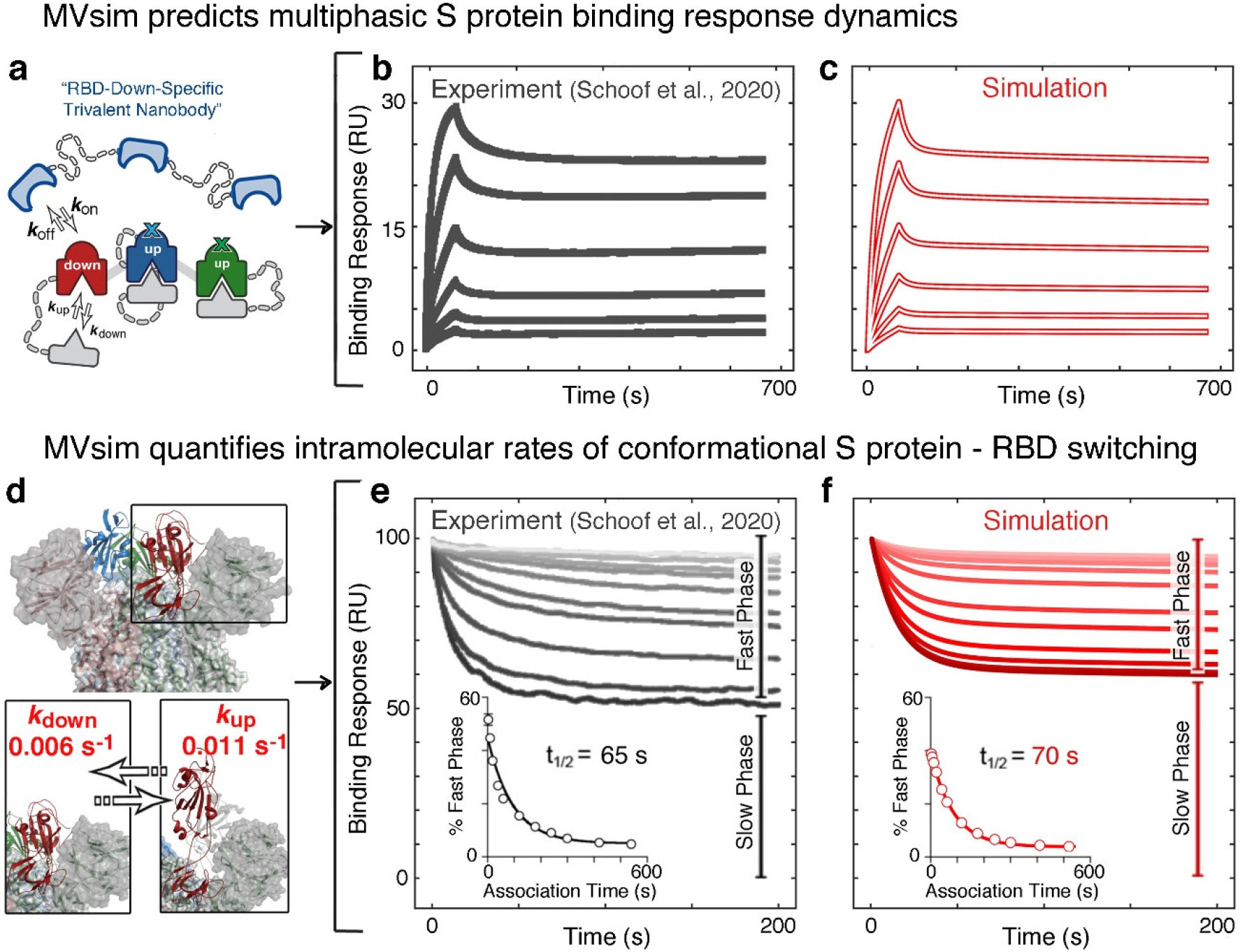
*MVsim* quantitatively predicts multiphasic binding response dynamics of S protein therapeutic neutralization and fits for rate constants of conformational switching. a. *MVsim* models the conformational change as an intramolecular ligand binding event (colored in grey) that toggles the trivalent S protein between “RBD-up” and “RBD-down” conformations. The conformational change is defined by a pair of first-order kinetic rate constants *k*_up_ and *k*_down_. **b,c,d**, The experimental kinetics of conformation-specific nanobody binding, adapted from Schoof et al.^37^, (**b**) are predicted by *MVsim* (**c**) and yield a single set of best-fit parameter values for *k*_up_ and *k*_down_ for an individual RBD (**d**). **e**, Conformational switching half-life experiments, also adapted from Schoof et al.^37^, alter the relative proportion of “slow phase” inhibitor dissociation (i.e., high avidity bivalent and trivalent interactions) and “fast phase” inhibitor dissociation (i.e., monovalent interactions) events. Here, due to relatively slow RBD switching rates, longer association times enable more S protein to be bound in high avidity interactions, and thus give rise to small percentages of “fast phase” dissociation events. **f**, To assess *MVsim* accuracy, the fitted parameters are used in a modeling framework to simulate the experimental system and compare half-lives (t_1/2_) of “RBD-up” S protein conformations.

The conformational dynamics of the S protein are of particular importance for understanding the mechanisms through which emerging SARS-CoV-2 mutational variants-of-concern (VOCs) increase infectiousness^45^. Mechanistically, S protein VOCs can function to stealth this protein from host immune surveillance, enhance the binding kinetics/affinity of the RBD-ACE2 interaction, stabilize the RBD-up configuration to increase the avidity of virion-host cell engagement, and/or augment conformational allostery that enables RBD binding to prime activation of membrane fusion^45^. To further apply *MVsim* to study S protein conformational dynamics, simulations were performed with the parameterized S protein RBD ensemble (Fig. S6a) to probe the effects of *k*_*up*_ and *k*_*down*_ on the ensemble of RBD configurations (Fig. S6b-d) and quantitatively relate these effects to ACE2 receptor binding on host cells (Fig. S6e-g). Here, with meaningful parameterization, *MVSim* may aid in mechanistically parsing the often multiple virulence-enhancing features that comprise most VOCs and to quantify, for example, the consequences of a mutational profile that simultaneously alters the RBD ensemble, enhances the kinetics of ACE2 binding, and reduces the potency of a neutralizing therapeutic.

## Discussion

*MVsim* is a new toolset created for the design, prediction, quantification, and mechanistic analysis of multivalent molecular interactions. It empowers users to simulate topologically complex multicomponent systems with an interactive GUI and to probe the relationships among configurational dynamics, cooperativity, effective concentration, and competitive binding that underlie the programmability of multivalent behavior. *MVsim* offers a considerable range of user inputs to parameterize the composition, kinetics, structure and topology, conformational flexibility, and component concentrations to simulate various disparate instances of multivalency in natural systems and synthetic designs.

Effectively simulating complex instances of multivalency has been hindered by the inherent combinatorial and spatial complexities that arise from binding domains sampling increasingly large, sterically constrained volumes to engage in a multitude of transient, pairwise interactions with unique energetic permissibilities^7,8,10^. *MVsim* addresses this challenge by combining rule-based modeling and multidimensional integrations to rapidly simulate system behavior by effectively tracking the evolution of hundreds of configurational binding state transitions throughout the lifetime of the molecular interaction. This generalized and extensible computational approach provided a means to create a consolidated modeling routine that yielded accurate and meaningful predictions of systems including simplistic beads-on-a-string topologies, intramolecular switches, and conformationally-regulated multicomponent assemblies.

Here specifically, we demonstrate the ability of *MVsim* to capture the unique multiphasic kinetics characteristic of multi-ligand, multispecific, and multivalent systems. We further show how *MVsim* can be used to predict and refine the design of systems that leverage multivalency to achieve nonlinear and ultrasensitive outputs, as well as the additional layering of multispecifity to create AND gate input/output operations. Further, we show use of *MVsim* in the advanced application of multispecificity toward the design of multivalent ligands capable of maximally distinguishing among pools of downstream targets and binding surfaces. Finally, to demonstrate the features and multiparameter inputs of *MVsim* applied to their fullest extent, a variety of therapeutically relevant structural features were computed for the SARS-CoV-2 S protein. Notably, *MVsim* was used to extract the effective concentration for the ACE2 interaction, quantifying the sterically unfavorable interaction that had previously been inferred from structural modeling, and to derive conformational rates of RBD switching dynamics from bulk kinetic measurements, previously quantifiable only though sophisticated single-molecule FRET experiments^46,47^.

The modular construction of *MVsim* also enables its straightforward extension to additional instances of multivalency. For example, additional configurational network tables can be applied to the source code to enable simulations of higher valencies and supramolecular topologies. Moreover, *MVsim* treats the calculation of effective concentration as an additional, modular mathematical step, enabling customization with any polymer end-to-end density function. Finally, the source code is further compatible with the MATLAB curve fitting toolbox to enable parameter estimation for incompletely characterized multivalent systems.

As presented here, *MVsim* simulates interactions between systems of receptors and ligands with valencies of up to three. The choice of trivalent interactions was chosen to balance the number of computational steps needed to map the complete configurational network, which scales factorially with valency, with the ability to model complex and important instances of multivalency, such as those that occur in bispecific antibodies and the trimeric architecture of the SARS-CoV-2 S protein. However, the configuration nomenclature, rule-based modeling, and combinatorial computation of effective concentrations that underlie the simulations are written in the source code to accommodate all valencies and numbers of competing ligands. Moreover, further approaches can be taken to overcome the computational demands of large systems of multivalent molecules. For example, additional rules can be added to the rule-based modeling routine to allow for sparse matrix sampling of the configurational microstate network and probability density functions to significantly decrease the computational time needed to simulate more topologically and combinatorially complex interactions. Nonetheless, there is a vast number of important biological questions and biomedical problems involving bivalent or trivalent molecular interactions, and *MVsim* provides tools to better analyze and engineer them.

## Methods

### MVsim

The *MVsim* multivalent simulation application was built within the MATLAB app development environment (version 2021a). Our previously reported microstate network model and odds-ratio-based calculation of effective concentration served as the foundation for creating a rule-based modeling routine for the enumeration of multivalent, multispecific, and multi-ligand receptor binding states^23^. Multidimensional integrations of the volumetric space sampled by hinged-linked binding domains were used to enable highly resolved determinations of effective concentration that are rapidly calculable across the nanoscopic and mesoscopic molecular scales relevant to complex multivalent systems.

### Experimental methods

Multivalent and multispecific receptors were constructed using the C-terminal SH3 domain of the human adaptor protein Gads and the synthetic protein Prb, as described previously^23^. Multivalent and multispecific ligands incorporated the SH3 binding peptide (SBP) from the Gads cognate ligand SLP-76^48^, as well as the synthetic designed Prb-binding DARPin, Pdar^49^. Association and dissociation kinetics between ligand and receptor constructs were quantified by surface plasmon resonance measurements on a Biacore S200 instrument.

### Documentation

A full set of version release notes, instructions, user tips, and annotated model source code are available on the *MVsim* homepage at https://sarkarlab.github.io/MVsim/.

## Supplementary Information

Extended methods, supplemental data and simulation figures, and *MVsim* source code are provided in the accompanying Supplementary Information.

## Acknowledgements

This work was supported by funding from the National Research Development and Innovation Fund (TKP2020 National Challenges Subprogram, Grant No. BME-NC) of the National Research Development and Innovation Office, the Higher Education Excellence Program (BME FIKP-BIO) of the Ministry of Human Capacities, and the Hungarian Scientific Research Fund (OTKA 119866) (to B.B.); from the National Institutes of Health (R35GM136309, R01GM113985, and R21EB022258 to C.A.S.); and from the Institute for Engineering in Medicine at the University of Minnesota (COVID-19 Rapid Response Grant to C.A.S.). The Biacore S200 instrument was made available through a shared instrument grant (S10OD021539) from the Office of Research Infrastructure Programs at the National Institutes of Health.

## Notes

https://sarkarlab.github.io/MVsim/

### Competing Interest Statement

The authors have declared no competing interest.

