## Supplementary Information for "*MVsim*: a toolset for quantifying and designing multivalent interactions"

##### 1. Extended Methods

##### 2. Supplemental Figures

Fig. S1: Expanded *MVsim* input features

Fig. S2: Expanded *MVsim* output features

Fig. S3: Mechanistic insights into multispecific and multi-ligand interactions through *MVsim* microstate analysis

Fig. S4: Design-build-test optimization of three-input AND gates

Fig. S5: Parameterization of *MVsim* to simulate the conformational dynamics, ACE2-RBD binding, and therapeutic neutralization of the SARS-CoV-2 S protein

Fig. S6: Simulating the effects of altered rate constants of RBD-switching on the S protein conformational ensemble and comparative binding to a neutralizing therapeutic or ACE2

##### 3. *MVsim* Source Code and User Documentation

**Extended Methods**

The basic framework of *MVsim* is based on state-space representation with gray-box modeling where the input and output of the model are measures proportional to the concentration of certain receptor-ligand conformations. *MVsim* generates the model structure based on rules determined by the experimental setup then estimates the model parameters based on simple experimental measurements and binding probability estimation.

### 1 Model structure

In a state-space representation of multivalent receptor (R)-ligand (L) binding, the state vector x(t) represents the concentrations of the different receptor-ligand conformations, and the state (or system) matrix **A**(t) contains the free ligand concentrations and rate constants. The output vector y(t) is a derived measure of the state concentrations. In surface plasmon resonance (SPR) experiments, y(t) is the product of matrix **C**, containing an appropriate SPR scaling constant times the molecular weights of the relevant states, and state vector x(t). Due to the inability to directly influence the state concentrations, there is no input vector in the model. This representation is equivalent to a system of kinetic reaction equations in vector-matrix form:

$$[R] \leftrightarrow -kon * [R] * [L] + koff * [RL]$$
$$[RL] \leftrightarrow +kon * [R] * [L] - koff * [RL]$$
$$\frac{d\begin{bmatrix} R \\ RL \end{bmatrix}}{dt} = \begin{bmatrix} -kon * L & +koff \\ +kon * L & -koff \end{bmatrix} \begin{bmatrix} R \\ RL \end{bmatrix}$$
$$\frac{dx(t)}{dt} = A(t) * x(t)$$
$$y(t) = C * x(t)$$

The corresponding MATLAB code representation of the kinetic equations generated based on the experimental setup using unique rate constant identifiers is as follows:

```
dydt = zeros(2,1);

dydt(1)=+Koffs.Koff_Receptor1_pos_1_Ligand1_pos_1*y(2)-
L1*Kons.Kon_Receptor1_pos_1_Ligand1_pos_1*y(1);

dydt(2)=+L1*Kons.Kon_Receptor1_pos_1_Ligand1_pos_1*y(1)-
Koffs.Koff_Receptor1_pos_1_Ligand1_pos_1*y(2);
```

#### 1.1 State vector

The state vector contains all possible conformations that can form from binding of receptor and ligand(s). The framework can model complex multivalent binding described by the following rules:

1) binding sites of the receptor(s)
2) binding sites of the ligand(s)
3) rules excluding binding between certain receptor-ligand binding sites
4) rules describing impossible conformations (e.g., due to steric blocking)

*MVsim* accommodates all models with the following rules: (1) one up to trivalent receptor, (2) up to three trivalent ligands, (3) receptor-site to ligand-site environment to specify exclusions between binding sites, and (4) allowing multiple binding events from the same type of ligand to the receptor.

These rules are phrased using unique identifiers of all the states from the perspective of a receptor without crosslinking. The identification system is as follows: the first three characters are the (1) identifier of the ligand, (2) the number of the ligand with a given identifier that is bound to the same receptor, and (3) the identifier of the ligand binding site on the first receptor binding site (e.g., Ligand A, first copy, second binding site is identified as A12 and the empty receptor is identified as 000). This is applied to all the receptor positions resulting in a 9-character long identifier (e.g., A11A12A13) in the case of trivalent receptors. Using these identifiers, all enumerated states comprise the state vector x(t).

#### 1.2 State matrix

The state matrix **A**(t) is populated following the logic that, if two states have the same identifiers except one of the triplets is 000 in one of the states, then the transition can happen (e.g., A11 000 A13 to A11 A12 A13). All of these transitions can be described with three constants:

1) If the transition is a dissociation (the number of 000s increases by one in the identifier), then the transition is described by a $k_{off}$ constant.
2) If the transition is an association (the number of 000s decreases by one in the identifier), then the transition is described by a $k_{on}$ constant multiplied by the ligand concentration.
    a. If the ligand is already bound to the receptor, this ligand concentration is an effective concentration estimated by the model.
    b. If the ligand is not yet bound to the receptor, this concentration is the bulk ligand concentration.

In SPR, both the ligand and effective concentrations are constant, but the model is capable of handling time-dependent changes in these concentrations if needed for the experimental setup. This is implemented in *MVsim* by changing the ligand concentrations at any point in the system.

The overall model structure of *MVsim* is based on the principles of chemical kinetics, and it does not introduce further biases or errors.

### 2 Model parameters

The model uses four types of parameters: association rate constant, $k_{on}$; dissociation rate constant, $k_{off}$; ligand concentration, [$L$]; and effective ligand concentrations, [$L_{eff}$].

The values for the association and dissociation rate constants are obtained from experiments with monovalent ligand and monovalent receptor (either in the literature or performed by the researcher). The ligand concentration is typically constant in a given phase of the experimental setup (nonzero during association and zero during dissociation), and SPR results are usually presented as a time-dependent plot of output (response units), often with an overlay of response curves from multiple ligand concentrations.

The last parameter is the effective ligand concentration, which can be crudely estimated from the radius of the receptor and ligand linkers, but which is computed more accurately by *MVsim* using a probabilistic approach that incorporates detailed biophysical properties of the interacting molecules.

#### 2.1 Estimation of effective ligand concentrations

From a probabilistic point of view, the effective ligand concentration is the concentration of a free monovalent ligand that has the same probability of binding to a free site on a partially bound receptor as the already partially bound multivalent ligand binding to the same free site on the same receptor.

To model the effective concentration, we use a probability density function (PDF) of the end-to-end distance, which in our case is the distance between the occupied binding site and the free binding site. PDF(r) for ligand and receptor describes the probability that the two ends are a distance r apart.

##### 2.1.1 Binding probability

For calculating a binding probability, we need PDF(x,y,z), a three-dimensional probability density function. The binding probability is the integral of the product of the ligand and receptor PDFs over the volume.

In the simplest case, the PDF is a uniform distribution of one molecule in a given volume. In the case of free ligand, we have a uniform distribution with a volume occupied on average by one molecule; in the case of a 1 M solution, the volume is $1/(6\times10^{23})$ $dm^3$.

*MVsim* uses more complex PDFs to model the receptor and ligand, based on literature on linker PDFs and an iterative process to calculate PDFs for joint linkers or linkers and spherical units. These derived PDFs can be used in a convolution integral to calculate the probability of binding and, by comparing it to that of the free ligand, we obtain effective concentrations.

###### *2.1.1.1 Ligand and receptor PDFs*

For the ligand and receptor PDFs, *MVsim* uses an iterative process which calculates the joint PDFs of linkers and binding domains. For example, a linear trivalent receptor molecule contains a binding domain, a linker, a second binding domain, a second linker, and a third binding domain. Therefore, a trivalent receptor is described by the joint PDF of these five components.

Since we are interested in the position of the two ends regardless of the positions of any intervening joining points of the components, we can integrate all the joining points leading to the following integral (see section 4.2 for additional details):

$$(f * g)(x_2, y_2, z_2) = \int_{-l}^{l}\int_{-l}^{l}\int_{-l}^{l} f\,(x, y, z) * g(x_2 - x, y_2 - y, y_2 - z)dxdydz$$

The iterative MATLAB implementation of the joint PDF is the following, where either a binding domain or a linker is added to the PDF:

```
fr = @(r0, fi0,r) f_root(r0).*f_toadd(sqrt(r.^2+r0.^2-
2.*r.*r0.*cos(fi0))).*r0.^2.*sin(fi0);
```

```
y(i) = quad2d(@(r0, fi0) fr(r0,fi0,x(i)) ,r0_min,r0_max,0, fi_max, 'AbsTol',
1e-10,'RelTol', 1e-4, 'MaxFunEvals', 20000);
```

###### 2.1.1.2 Calculating effective ligand concentration from binding probability

The equation for converting the multivalent binding probability to an effective concentration is as follows (see section 4.1 for details):

$$C_{eff} = \frac{\int PDF_{ligand}(V) * PDF_{receptor}(V)dV}{constant_{PDF\ normalisation}}$$

which is implemented in *MVsim* with the following code:

```
final_conv =  @(r,z)
f_Receptor(sqrt(r.^2+z.^2)).*f_Ligand(sqrt(r.^2+(z+shift).^2)).*r;

value = 2*pi*integral2(final_conv, 0,min(length_Receptor, length_Ligand), -
min(length_Receptor, length_Ligand,min(length_Receptor, length_Ligand);

Ceff = value/(c_norm_Receptor*c_norm_Ligand)/(6*(10^-4));
```

###### 2.1.1.3 Solving integrals for the binding probability

To determine binding probabilities in *MVsim*, we face the practical problem of computing triple integrals with complex or arbitrary functions, which are infeasible to solve over a reasonable time period. We therefore rewrite all integrals in polar coordinates using the spherically symmetric nature of the PDF, resulting in more tractable double integrals (see sections 4.3 and 4.4 for details).

### 3 Output

The y(t) output vector and the **C** output matrix convert the microstate concentrations to the desired output format.

In the case of SPR, y(t) has only one element, the output in RU, which is proportional to the molecular weight(s) of the bound ligands. To get this output, C is a one-dimensional matrix with the sum of the molecular weights of the ligands bound to the receptor times the volume of the SPR chip times the conversion constant from grams to picograms.

*MVsim* also provides additional output options. The output can be in the form of molar concentration or weight/RU and y(t) can contain all the states or the sum of states based on their binding valency or the type of the bound ligands.

The model outputs are presented in conventional time-dependent plots as well as in a graphical representation of the microstates that sheds light on the species driving noncanonical binding dynamics in multivalent reactions.

### 4 Mathematical details of the model

#### 4.1 Calculating effective concentrations

The effective molecule number $(Eff_{num})$ is the number of monovalent ligands that are equally probable to bind a free binding domain on a multivalent receptor as the probability of an already-receptor-bound multivalent ligand binding with another one of its domains to the same free binding domain on the receptor:

$$Eff_{num} * P(uniform\ ligand\ position = receptor\ free\ end\ position)$$
$$= P(ligand\ free\ end\ position = receptor\ free\ end\ position)$$

$$Eff_{num} = \frac{P(ligand\ free\ end\ position = receptor\ free\ end\ position)}{P(uniform\ ligand\ position = receptor\ free\ end\ position)}$$

To get the probability of the two ends meeting, we can examine all points in space, note the probability of the ends meeting at each point, and integrate over all these values to get the overall probability in question.

$$P(ligand\ free\ end\ position = receptor\ free\ end\ position)$$
$$= \int PDF_{ligand}(V) * PDF_{receptor}(V)dV$$

and

$$P(uniform\ ligand\ position = receptor\ free\ end\ position)$$
$$= \int 1 * constant_{PDF\ normalisation} * PDF_{receptor}(V)dV$$
$$= constant_{PDF\ normalisation} * \int PDF_{receptor}(V)dV = constant_{PDF\ normalisation}$$

Therefore the effective number of ligands can be calculated by:

$$Eff_{num} = \frac{\int PDF_{ligand}(V) * PDF_{receptor}(V)dV}{constant_{PDF\ normalisation}}$$

##### 4.1.1 Effective ligand concentration

We can convert the effective number of ligands to an effective ligand concentration by dividing this ligand number by Avogadro's number and the accessible volume.

#### 4.2 PDF for linkers and binding domains

##### 4.2.1 Linker PDF

The PDF for the ligand end-to-end distance in terms of contour length ($L_c$) and persistence length ($L_p$) is

$$PDF(r, L_p, L_c) \propto \frac{1}{\left(1-\frac{r^2}{L_c^2}\right)^{\frac{9}{2}}} e^{\left(-\frac{9L_c}{8L_p}\frac{1}{1-\frac{r^2}{L_c^2}}\right)}$$

or

$$PDF(x, y, z, L_p, L_c) \propto \frac{1}{\left(1-\frac{x^2+y^2+z^2}{L_c^2}\right)^{\frac{9}{2}}} e^{\left(-\frac{9L_c}{8L_p}\frac{1}{1-\frac{x^2+y^2+z^2}{L_c^2}}\right)}$$

##### 4.2.2 Binding domain PDF

The PDF of the binding domain is based on its diameter (d) as follows:

$$PDF(r, d) \propto \{if\ d-\epsilon \leq r \leq d+\epsilon = 1, otherwise = 0\}$$

or

$$PDF(x, y, z, d) \propto \{if (d-\epsilon)^2 \leq x^2+y^2+z^2 \leq (d+\epsilon)^2 = 1, otherwise = 0\}$$

where $\epsilon$ represents the uncertainty related to the domain diameter (e.g., standard deviation of its measurement).

#### 4.3 Convolution of two PDFs

The base equation of convolution is:

$$(f * g)(t) = \int_{-\infty}^{\infty} f(\tau) g(t-\tau)\, d\tau$$

The same equation using (x,y,z) coordinates and starting from a non-zero point is:

$$(f * g)(x_0, y_0, y_0, x_2, y_2, z_2) = \int_{-l}^{l}\int_{-l}^{l}\int_{-l}^{l} f(x-x_0, y-y_0, z-z_0) * g(x_2-x, y_2-y, z_2-z) dxdydz$$

If $x_0 = 0, y_0 = 0, z_0 = 0$, this reduces to:

$$(f * g)(x_2, y_2, z_2) = \int_{-l}^{l}\int_{-l}^{l}\int_{-l}^{l} f(x, y, z) * g(x_2-x, y_2-y, y_2-z) dxdydz$$

And switching to spherical coordinates:

$$(f * g)(R \mid r_0) = \int_0^{2\pi}\int_0^{\pi}\int_0^{r_{max}} f(\| \hat{r}-\widehat{r_0} \|) * g\left(\| \hat{R}-\hat{r} \|\right) r^2 sin\phi drd\phi d\theta$$

$$x = r\, sin(\phi) cos(\theta), y = rsin(\phi) sin(\theta), z = rcos(\phi)$$

$\parallel \hat{R} - \hat{r} \parallel$
$= \sqrt{R^2 + r^2 - 2Rr[sin(\phi_R)sin(\phi)cos(\theta_R)cos(\theta) + sin(\phi_R)sin(\phi)sin(\theta_R)sin(\theta) + cos(\phi_R)cos(\phi)]}$

Setting $\phi_R$ to zero, we get:

$$\parallel \hat{R} - \hat{r} \parallel = \sqrt{R^2 + r^2 - 2Rr[0 * sin(\phi)cos(\theta_R)cos(\theta) + 0 * sin(\phi)sin(\theta_R)sin(\theta) + 1 * cos(\phi)]}$$

$$\Longrightarrow \parallel \hat{R} - \hat{r} \parallel = \sqrt{R^2 + r^2 - 2Rrcos(\phi)}$$

where r is the linker end-to-end distance, R – r is the hinge-to-binding site distance in the domain, and R is the point where we are interested in the probability of the PDF:

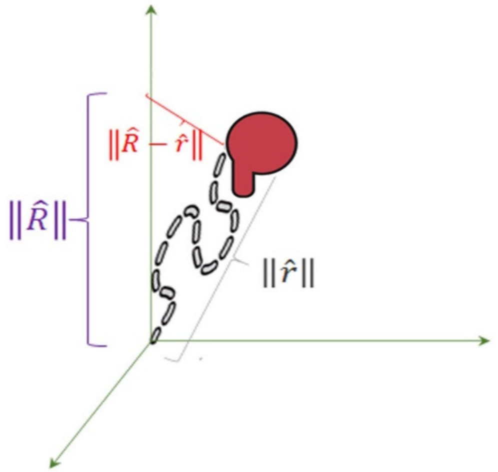


If $r_0$ is zero and we position R to fall to the z-axis, then $\phi_R$ is zero, and we get the following function for PDF(R):

$$(f * g)(R) = \int_0^{2\pi} \int_0^{\pi} \int_0^{r_{max}} f\,(r) * g\left(\sqrt{R^2 + r^2 - 2Rrcos\phi}\right) r^2 sin\phi drd\phi d\theta$$

$$(f * g)(R) = \int_0^{2\pi} 1\,d\theta * \int_0^{\pi} \int_0^{r_{max}} f\,(r) * g\left(\sqrt{R^2 + r^2 - 2Rrcos\phi}\right) r^2 sin\phi drd\phi$$

$$(f * g)(R) = 2\pi \int_0^{\pi} \int_0^{r_{max}} f\,(r) * g\left(\sqrt{R^2 + r^2 - 2Rrcos\phi}\right) r^2 sin\phi drd\phi$$

#### 4.4 Convolution with actual functions

Let us consider a domain-linker-domain construct starting at the origin ($r_0$ = 0), with the first domain ending at distance $r_1$, the linker ending at distance $r_2$, and the second domain ending at distance $r_3$. We can further define R as the endpoint of the entire construct of interest (here, equal to $r_3$) and also define r as any intermediate endpoint of interest (here, $r_1$ or $r_2$).

The convolution of two functions corresponding to the first domain and the linker is therefore given by:

$$\left(f_{domain,1} * g_{linker}\right)(r_2 \mid r_0 = 0) = 2\pi \int_0^{\pi} \int_0^{r_{max}} f_{domain,1}(r_1) * g_{linker}\left(\sqrt{r_2^2 + r_1^2 - 2r_2 r_1 cos\phi_1}\right) r_1^2 sin\phi_1 dr_1 d\phi_1$$

And the convolution of three functions, corresponding to the full domain-linker-domain construct, is:

$$\left(f_{domain,1} * g_{linker} * f_{domain,2}\right)(R \mid r_0 = 0) = 2\pi \int_0^{\pi} \int_0^{r_{max}} \left(2\pi \int_0^{\pi} \int_0^{r_{max}} f_{domain,1}(r_1) * g_{linker}\left(\sqrt{r_2^2 + r_1^2 - 2r_2 r_1 cos\phi_1}\right) r_1^2 sin\phi_1 dr_1 d\phi_1\right) * f_{domain,2}\left(\sqrt{R^2 + r_2^2 - 2Rr_2 cos\phi_2}\right) r_2^2 sin\phi_2 dr_2 d\phi_2$$

Since the two-function convolution only depends on one parameter, $r_2$, the value of this convolution can be fit to a new function, $f_{root}$, to reduce the three-function convolution to a two-function convolution of $f_{root}$ and the remaining function, $f_{to\_add}$. In the present example, this would be written as:

$$\left(f_{domain,1} * g_{linker} * f_{domain,2}\right)(R \mid r_0 = 0) = 2\pi \int_0^{\pi} \int_0^{r_{max}} f_{root}(r_2) * f_{to\_add}\left(\| \hat{R} - \hat{r_2} \|\right) r_2^2 sin\phi dr_2 d\phi$$

where $f_{root} = \left(f_{domain,1} * g_{linker}\right)(r \mid r_0 = 0)$ and $f_{to\_add} = f_{domain,2}(\| \hat{R} - \hat{r} \|)$.

This is a more computationally efficient double integral and thus allows for faster calculation of convolution chains.

#### 4.5 Effective ligand number with predicted PDFs

For calculating effective ligand number, we need the free-end to bound-end PDFs for the ligand and the receptor, and we can predict their non-normalized PDFs using the technique described in section 4.4.

Therefore, the equation for the effective number of ligands is:

$$Eff_{num} = \frac{\int C_L * f_{ligand}(V) * C_R * g_{receptor}(V) dV}{C_U}$$

where $C_U$, $C_L$, $C_R$ are normalization constants for the uniformly distributed ligand, the bound ligand, and the bound receptor, respectively. And,

$$f_{ligand}(V) = L\left(\left(f_{L,domain,i} * g_{L,linker,i}\right)_{m-1} * f_{L,domain,m}\right)(r_m, \phi, \theta)$$

$$g_{receptor}(V) = R\left(\left(f_{R,domain,i} * g_{R,linker,i}\right)_{n-1} * f_{R,domain,n}\right)(r_n, \phi, \theta)$$

where m and n are the valencies of the ligand and receptor, respectively and $L(r_m, \phi, \theta)$ and

$R(r_n, \phi, \theta)$ are the entire ligand and receptor PDFs. The rank of the domains and linkers is i = 1 to m for the ligand and i = 1 to n for the receptor.

Together, this yields the equation

$$Eff_{num} = \frac{4\pi C_L C_R \int R\left(\left(f_{R,domain,i} * g_{R,linker,i}\right)_{n-1} * f_{R,domain,n}\right)(r_n = r) * L\left(\left(f_{L,domain,i} * g_{L,linker,i}\right)_{m-1} * f_{L,domain,m}\right)(r_m = r) r^2 dr}{C_U}$$

#### 4.6 Further technical considerations

##### 4.6.1 Integral bounds

Matching the integral bounds of a convolution of $f_{root}$ and $f_{to\_add}$ to the input ranges of these functions significantly speeds up numerical integration. For PDF calculations using spherical integral convolution, the optimized bounds are:

$\theta$: $[0, 2\pi]$ (in our case, it integrates to $2\pi$)

$\phi$: if $R > L_{root}$, then the bounds are $\left[0, arcsin\left(\frac{L_{to\_add}}{R}\right)\right]$; otherwise, $[0, \pi]$

r: $\left[max\left(0, R - L_{to\_add}\right), min(L_{root}, R + L_{to\_add})\right]$

where $L_{root}$ is the maximum of the input range of the first function ($f_{root}$), $L_{to\_add}$ is the maximum of the input range of the second function ($f_{to\_add}$), and R is the position at which we want to evaluate the integrals.

If a binding domain is added, the bounds of r are $[d - \epsilon, d + \epsilon]$.

##### 4.6.2 Flipping the origin

Due to the commutative nature of the convolution, the following applies:

$$(f * g)(t) := \int_{-\infty}^{\infty} f(\tau) g(t - \tau)\, d\tau = \int_{-\infty}^{\infty} f(t - \tau) g(t)\, d\tau$$

Numerically integrating the convolution generally leads to rapid calculations. However, under certain conditions (e.g., when the domains of the two functions are of different orders of magnitude and thus large regions of the sampled space have zero values), it takes significant time for the algorithm to converge. In these cases, it is beneficial to use the function with the smaller coordinate domain as $f(\tau)$. It does not change the outcome, but can greatly reduce the computational time.

##### 4.6.3 Shift

For the final convolution, the PDFs of the ligand and the receptor can start from the same origin, the center of the binding site. However, in cases where two domains are not binding in an inline configuration but in a twisted configuration, it is computationally useful to introduce a spatial shift to compute the final convolution since the twisted binding configuration introduces complex angular restrictions due to the shifted relative positioning of the domain-linker hinges. The magnitude of the shift parameter is determined by the diameters of the receptor and ligand binding domains. In these cases, the convolution changes from $Eff_{num} = \frac{C_L * C_R \int\int\int R(r) * L(r) r^2 \sin(\phi) dr d\phi d\theta}{C_U}$ to $Eff_{num} = \frac{C_L * C_R * 4\pi \int R(r) * L(r) r^2 dr}{C_U}$ in the case of no applied shift or to $Eff_{num} = \frac{C_L * C_R \int\int\int R(r) * L\left(\sqrt{shift^2 + r^2 - 2shift * r cos\phi}\right) r^2 sin(\phi) dr d\phi d\theta}{C_U}$ in the case of an applied shift. This translational shift is more easily and quickly applied in cylindrical coordinates along the z axis: $Eff_{num} = \frac{C_L * C_R * 2 * \pi \int\int R(\sqrt{r^2 + z^2}) * L(\sqrt{r^2 + (z + shift)^2} * r \, dr \, dz}{C_U}$.

#### Supplemental Figures

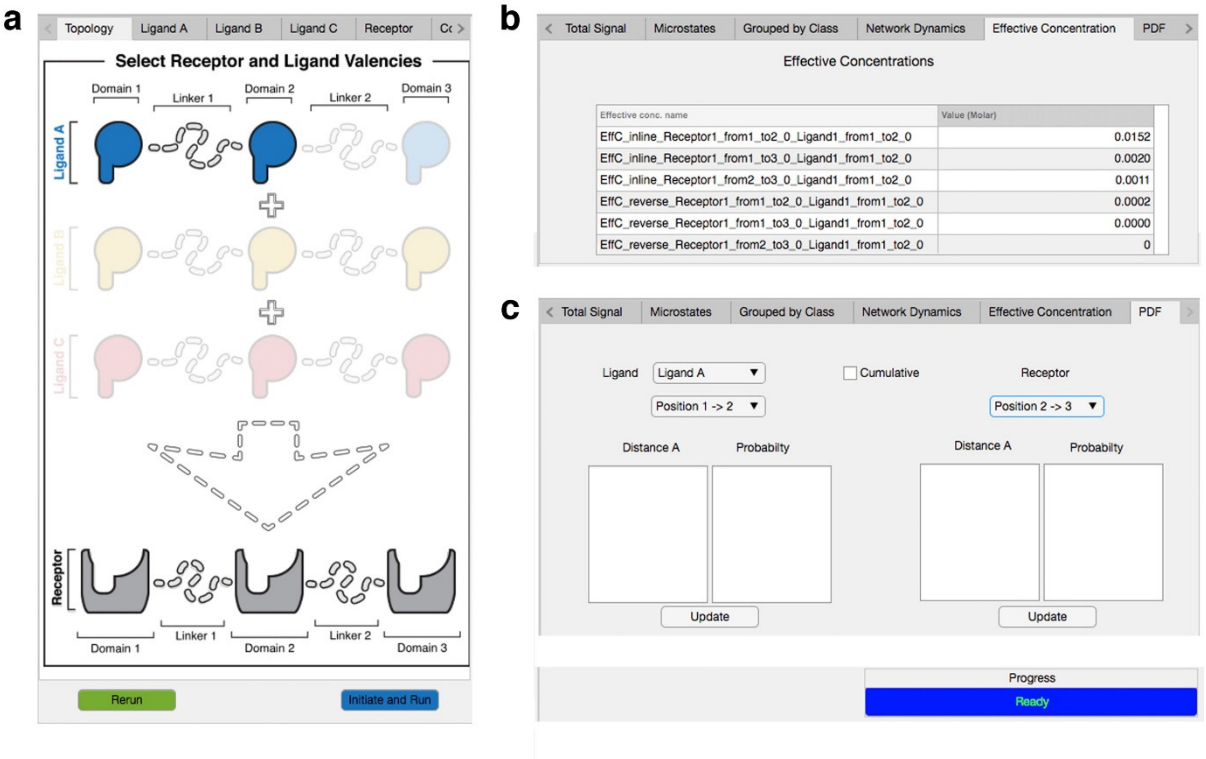


**Fig. S1 [related to Fig. 1]: GUI Inputs. *MVsim* enables bypassing of the probability density and effective concentration calculations by directly specifying either a set of effective concentrations, [$L_{eff}$], or by uploading a set of probability density functions (PDFs) for a multivalent system of interest. a**, An example of a multivalent system. Here, a bivalent ligand (blue) interacts with a trivalent receptor (grey). **b**, After initiating the simulation, and upon its completion, the "Effective Concentration" tab in the output window displays the name of each calculated [$L_{eff}$] (left column) and its calculated value in units of molar concentration (right column). The user can edit each [$L_{eff}$] value by clicking on corresponding field in the right column. The user-specified [$L_{eff}$] values are computed through use of the "Rerun" button (colored green; **a**, bottom left). **c**, The "PDF" tab in the output window features an interface through which PDFs can be crafted and applied to the user-specified multivalent system; this can be performed for any or all of the ligand(s) and the receptor, as well as for any or all of the [$L_{eff}$] values. To begin, the user selects the multivalent molecule they wish to edit from the "Ligand" drop-down menu. Next, for ligand and/or receptor, the user selects the positional distribution they wish to edit from the lower drop-down menu. Here, for example, the selection of "Ligand A" and "Position 1 $\rightarrow$ 2" allows the user to edit the PDF that describes the spatial distribution of the second binding domain relative to the first within the trivalent structure of Ligand A. Coordinates for the PDF are entered in the "Distance" (x-axis values measured in Angstroms) and "Probability" fields and are uploaded in place of the existing calculated values using the "Update" button. To perform simulations with updated PDFs, the "Rerun" button is used.

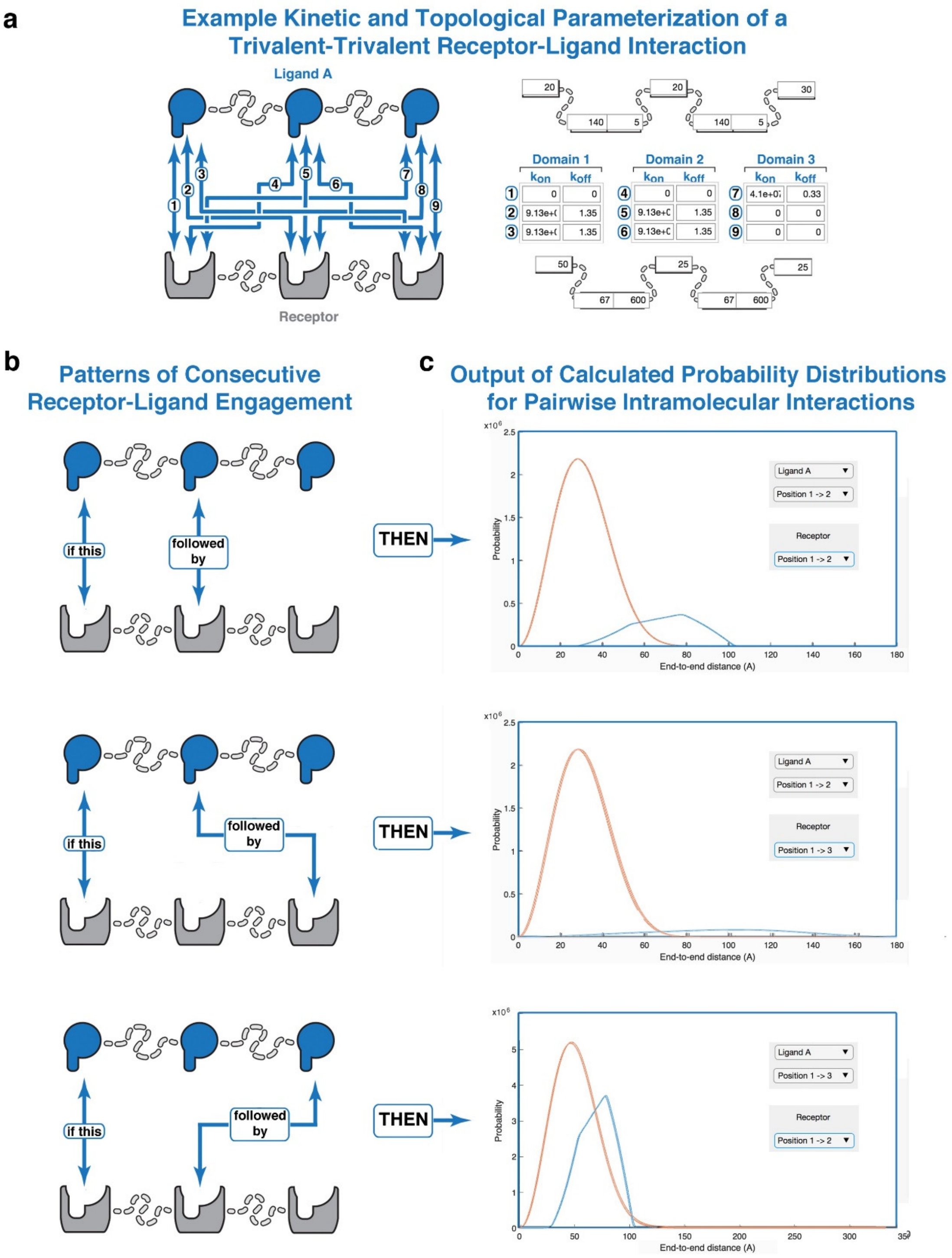


**Fig. S2 [related to Fig. 2]: GUI Outputs. *MVsim* enables visualization of the computed set of probability density functions (PDFs) for a specified multivalent system. a**, As an example, a trivalent-trivalent receptor-ligand interaction is selected (left panel) and parameterized with a set of kinetic rate constants and topological parameters (right panel). **b**, Effective ligand concentrations [$L_{eff}$] are derived from the spatial proximity between ligand and receptor binding domains that results from the tethering via a prior point of interaction. Here, three [$L_{eff}$] possibilities are shown. In each of the three examples, the initial "if this" interaction occurs between the first binding domain of the ligand and the first binding domain of the receptor. Subsequently, three of the four possible "followed by" binding events are depicted. **c**, For each of these three [$L_{eff}$] possibilities, one-dimensional representations of the PDFs are shown for each, consisting of both the domain-to-domain distributions for the ligand (orange PDF) and receptor (blue PDF).

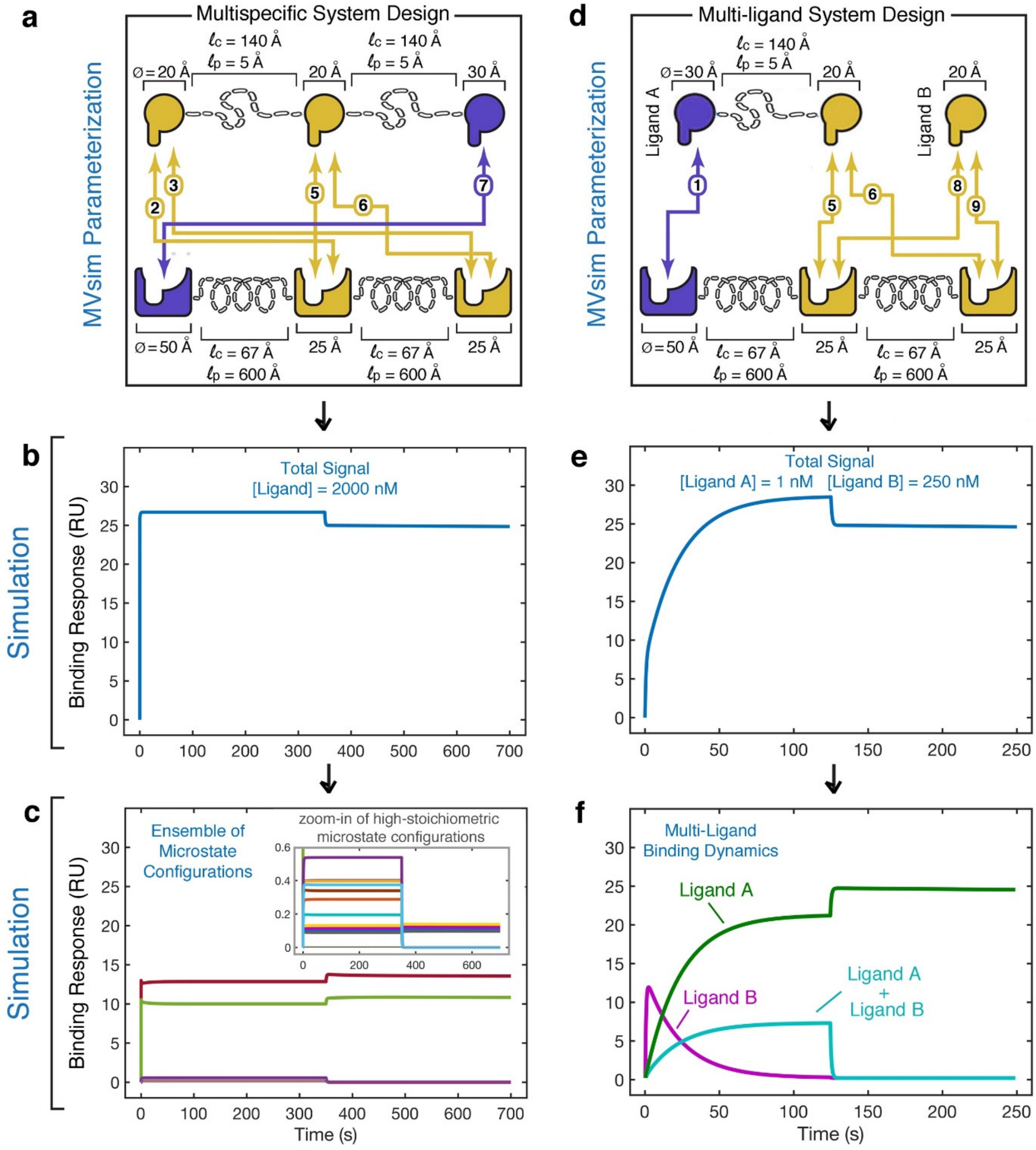


**Fig. S3 [related to Fig. 3]: *MVsim* provides mechanistic insights into multivalent interactions by enabling visualization of the underlying configurational microstate dynamics.** **a**, *MVsim* parameterization used to predict and simulate the experimental multispecific binding response dynamics shown in Fig. 3e. **b**, Total signal from the simulated response dynamics with a ligand concentration of 2000 nM. **c**, The ensemble of microstates for the trace in (**b**) shows the respective contributions of the two trivalent configurations (colored dark red and green, in main plot) and the multitude of high-stoichiometric, mixed monovalent-bivalent configurations (zoomed inset) to the overall multiphasic binding response dynamics. **d**, *MVsim* parameterization used to predict and simulate the experimental multi-ligand binding response dynamics shown in Fig. 3h. **e**, Total signal from the simulated response dynamics with a Ligand A concentration of 1 nM and a Ligand B concentration of 250 nM. **f**, The respective ligand-bound states for the trace in (**e**) shows the respective contributions of the bivalent, higher-avidity/slower-associating Ligand A and fast on/off binding of the monovalent Ligand B towards the multiphasic binding response dynamics.

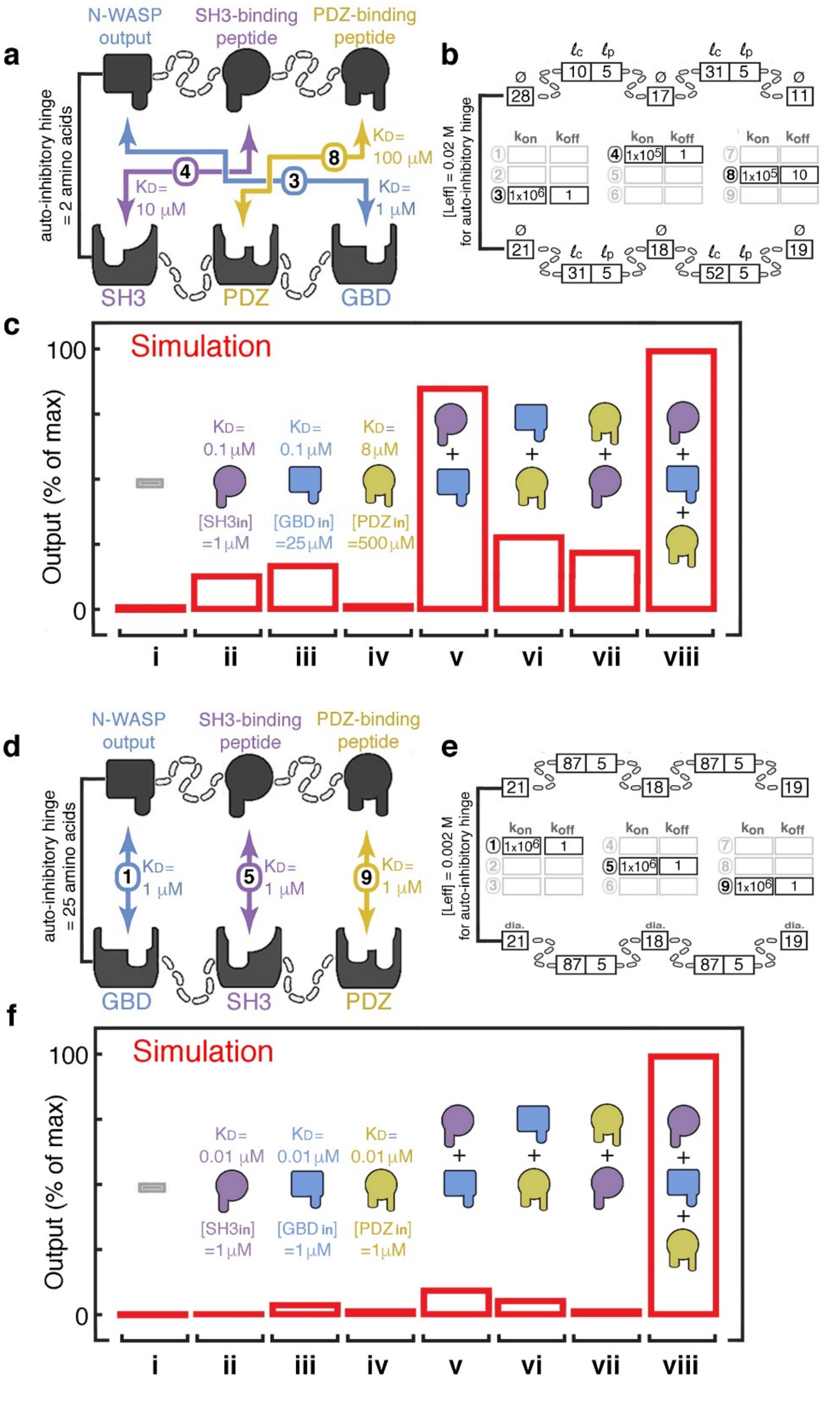


**Fig. S4 [related to Fig. 4]: *MVsim* guides the design, building, and testing of molecular logic gates for use in synthetic biology. a**,**b**, The architecture and experiment-based parameterization of the three-input AND gate simulated in Fig. 4b. **c**, Simulated performance of the AND gate detailed in (**a**,**b**) in the presence of (from left to right) i. No Input; ii, iii, iv. Single inputs; v, vi, vii. Dual inputs; and viii. Triple input. Significant spurious activation of the AND gate's output is predicted for both single and dual inputs, particularly the dual SH3 (colored purple) and GBD (colored blue) inputs in (v.). **d**,**e**, Using *MVsim* to sample the parameter space of a three-input AND gate identifies and optimized theoretical construction that avoids contorted and unfavorable binding configurations present in (**a**) through an inline topology (**d**), longer linkages and matched set of kinetics rate constants of association and dissociation (**e**). **f**, Simulated performance of the optimized, theoretical three-input design shows suppressed spurious activation in the presence of single and dual inputs.

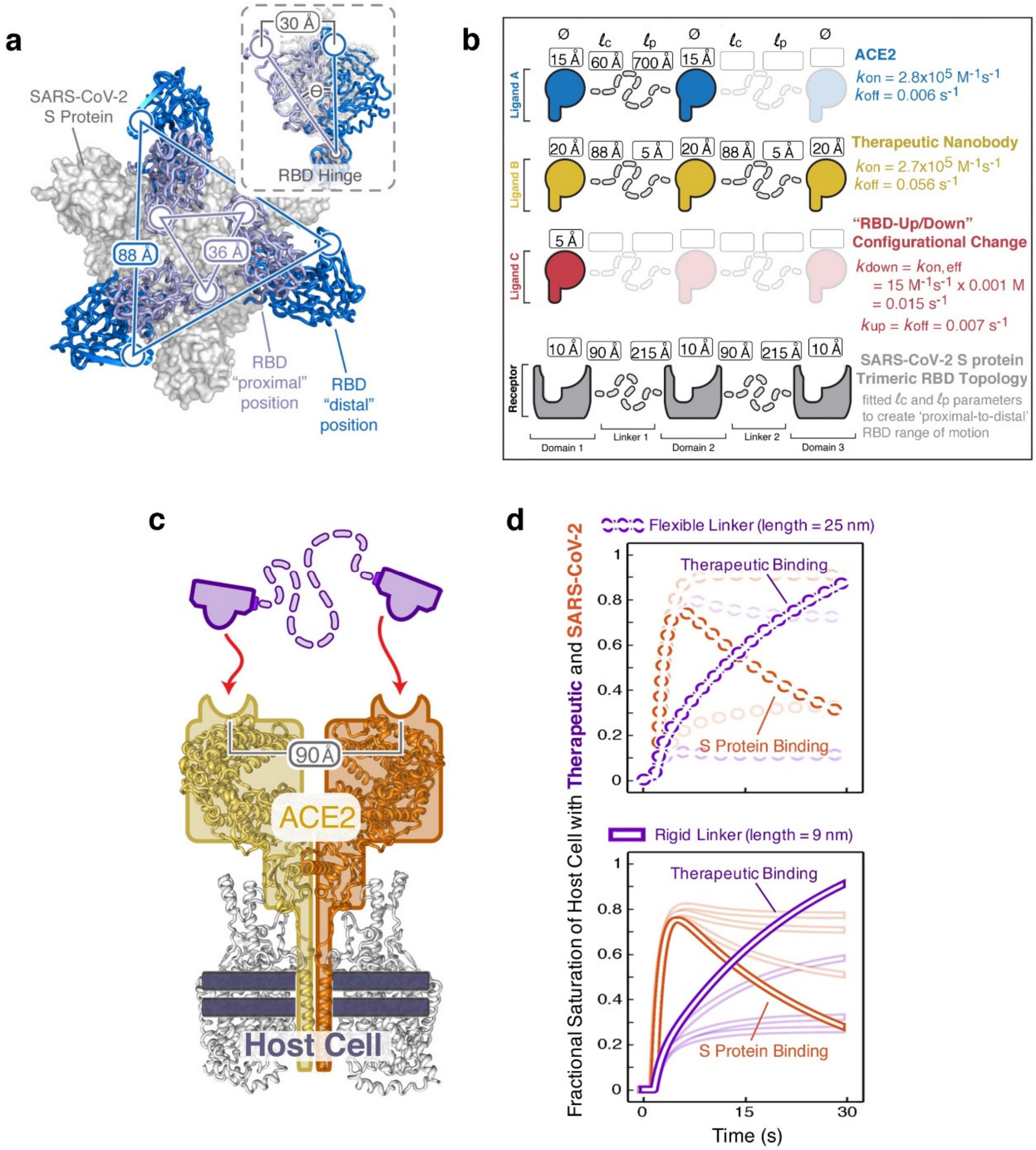


**Fig. S5 [related to Fig. 5]: *MVsim* can predict SARS-CoV-2 S protein binding response dynamics through parameterization with a combination of experimentally-determined parameters and fitted parameters extracted from structural and binding studies. a**, Superposition of 10 cryo-EM structures show a span of conformations that the three S protein receptor binding domains (RBDs) can sample. The limits of motion captured in these static structures can be used to parameterize *MVsim*. **b**, A set of topological and kinetic parameters (domain diameter, $\phi$; linker contour length $\ell$c; and linker persistence length, $\ell$p) were derived that can approximate the constraints of the S protein (grey), ACE2 (blue), and RBD configurational dynamics (red) using the linear, beads-on-a-string topology that *MVsim* is based upon. These parameter values were used for the simulations in Fig. 5. **c**, In addition to the strategic therapeutic targeting of the S protein (as explored in Fig. 5c-f), binders targeted to the RBD-binding surface of the ACE2 dimer have been assessed for their therapeutic potential. Here, the available structures for full length ACE2 and the ACE2-RBD complex were used to parameterize *MVsim* with descriptive multivalent topologies, domain diameters, kinetic rate constants of association and dissociation, and contour and persistence lengths that provide bounds for the captured ranges of motion. Simulations were performed to determine the optimal lengths for both flexible linkages (**d**, top plot) and rigid linkages (**d**, bottom plot) that are able to provide maximal protective shielding (darker purple curves in both plots) and therefore result in the largest reductions of SARS-CoV-2 infection (darker orange curves in both plots). *MVsim* predicts that level of protection that a shielding therapeutic can provide when the linker connecting the two binding regions is changed.

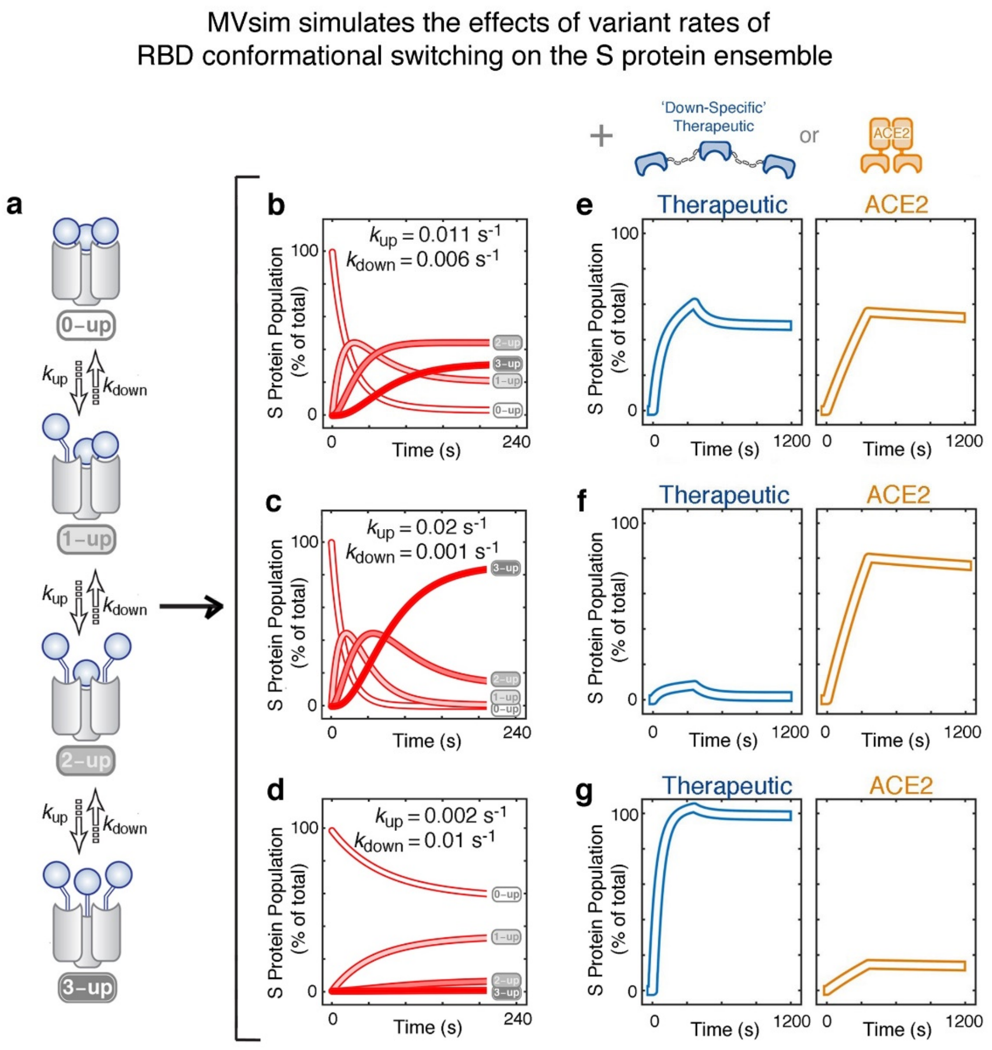


**Fig. S6 [related to Fig. 6]: *MVsim* enables the parameterized SARS-CoV-2 S protein simulation to model the consequences of variation of the RBD-up/down rates of switching on the S protein conformational ensemble.** **a**, *MVsim* can simulate the conformational ensemble of S protein RBD states as a function of changing RBD rate constants (e.g., those hypothesized to occur in some S protein mutational variants). **b**,**c**,**d**, *MVsim* modeling of the system with $k_{up}$:$k_{down}$ ratios of (**b**) 1.8, (**c**) 20, and (**d**) 0.2 show simulated effects on the ensemble of four RBD configurations (0-up, 1-up, 2-up, and 3-up). **e**,**f**,**g**, To further simulate the effects of variant rates of RBD conformational switching, the simulated S proteins (**b**,**c**,**d**) were each introduced to either a "down-specific" trivalent therapeutic nanobody (as treated in Fig. 6) or a dimeric ACE2. Therapeutic and ACE2 binding were performed as simulated SPR experiments with an association phase (0-300 s) and a dissociation phase (300-1200 s).
